# Microbes as part of ancestral neuronal circuits: Bacterial produced signals affect neurons controlling eating behavior in *Hydra*

**DOI:** 10.1101/2023.04.28.538719

**Authors:** Christoph Giez, Denis Pinkle, Yan Giencke, Jörg Wittlieb, Eva Herbst, Tobias Spratte, Tim Lachnit, Alexander Klimovich, Christine Selhuber-Unkel, Thomas Bosch

## Abstract

Although recent studies indicate the impact of microbes on the central nervous systems and behavior, it remains unclear how the relationship between the functionality of the nervous system, behavior and the microbiota arise. We studied the eating behavior of Hydra, a host that has a simple nervous system and a low-complexity microbiota. To identify the neuronal subpopulations involved, we used a subpopulation specific cell ablation system and calcium imaging. The role of the microbiota was uncovered by reducing the diversity of the natural microbiota. Here, we demonstrate that different neuronal subpopulations are functioning together to control the eating behavior. The microbiota participates in control of the eating behavior since germ-free or mono-colonized animals have drastic difficulties in mouth opening. This was restored by adding a full complement of the microbiota. In summary, we provide a mechanistic explanation of how the eating behavior is controlled in *Hydra* and how microbes can affect the neuronal circuit.

**Highlights:**

- Multiple neuronal modules and their networks control complex behavior in an animal lacking a central nervous system.
- Its associated microbes participate in these neuronal circuits and influence the eating behavior.
- Disorganization of the microbiota negatively impacts this eating behavior.
- Glutamate participates in an evolutionary ancient interkingdom language.

## Introduction

Understanding the neuronal basis of any behavior is a challenging task, given the complex interplay between neuronal circuits and the internal state of the organism such as hunger, fear or motivation^1,2^. The neuronal basis for behavior has been studied in various animal hosts. In *Drosophila* larvae, food deprivation shapes the olfactory behavior, highlighting the role of state-dependent neuronal circuits in dynamic behaviors^3^. Another example is the mating and aggression behavior in mice and flies, where a small number of neurons appear to control both behaviors in a state-dependent manner^4^. To add another level of complexity, gut microbes can affect behavior as well as activity of the nervous system^5–9^. For instance, the gut microbiota can affect aggression, fear, motivation, hunger, and emotional behaviors^10–18^. However, how microbes actively regulate the nervous system and thereby affect internal states and behaviors remains mostly unknown. It can be expected that the myriad of neurochemicals produced by microbes that live in close association with their host can influence neuronal activity. For example, muropeptides produced by the gut microbiota of mice are sensed by neurons expressing the Nod2 receptor in a specific region of the brain, which affects feeding behavior^19^.

Ideally, the complex interaction between behavior, internal state and microbes should be studied in a host that displays complex behavioral patterns, but also has a simple nervous system and a microbiota that is not too complex. Such a host is *Hydra* (Fig. 1A), a member of the phylum Cnidaria, which forms the sister group to the Bilateria in the Eumetazoa clade, which includes one of the earliest animals with a nervous system (Fig. 1B) ^20^. The nervous system of *Hydra* is composed of only two main neuronal cell types, sensory and ganglion cells, that form a nerve net in the ectoderm and endoderm, respectively^21–23^. The associated microbiota colonizes the glycocalyx that overlays the ectoderm and has direct contact to ectodermal sensory cells (Fig. 1C). There is no cephalization or ganglion formation, but regions along the body column have higher or lower densities of specific neuronal subpopulations (Fig. 1D)^24,25^. These represent non-overlapping neuronal networks with differential activity^26^. Already in 1744, Abraham Trembley recognized that *Hydra*’s behavior can be either spontaneous (such as contractions) or stimuli-evoked^27^. An example of the latter is the eating behavior, schematically shown in Fig. 1E, that can be induced by food or food-associated molecules such as reduced glutathione (GSH)^28,29^. The pattern is stereotypical and consists of three different stages: tentacle writhing, tentacle ball formation, and mouth opening^30^ (Fig. 1E, F, Suppl. Video 1). In animals without nerve cells this behavior is completely absent^31^. Previous work had indicated that neurons in the head region and not in the body column or tentacles were involved in the eating behavior^32,33^. More recently, single cell RNA sequencing had identified different neuronal subpopulations including those in the head^24,25,34^. This elaborate behavior also responds to an internal state, since a food stimulus given to well-fed animals does not result in a complete eating behavior response^28,29,35^.

**Figure 1.**
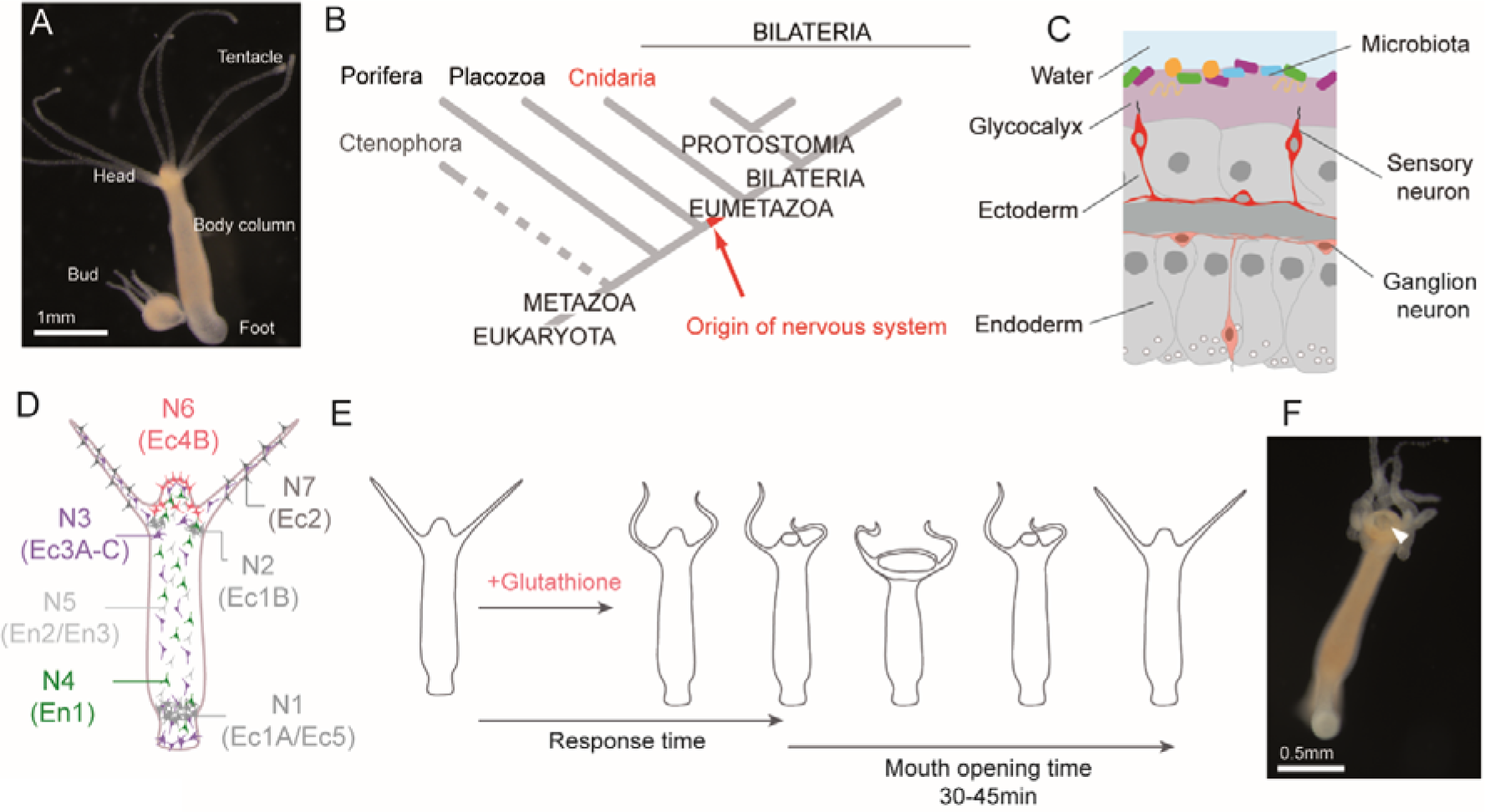
*Hydra vulgaris* AEP as a model system to study neuro-bacteria interactions. **A.** *Hydra* polyp with a forming bud (asexual reproduction). Scale 1mm. **B.** Phylogenetic position of *Hydra* in the phylum of Cnidaria which is the sister group of the Bilateria. **C.** Tissue organization and localization of microbiota on the glycocalyx at the outside of the polyp. **D.** Schematic presentation of the seven neuronal subpopulations and their distribution (after Siebert *et al.* 2019 and Klimovich *et al*. 2020)^24,25^. The alternative nomenclature includes Ec for ectoderm and En for endoderm. The head region contains N6, N3 and N4 neurons. **E.** The eating behavior of *Hydra* towards glutathione as defined by Loomis and Lenhoff ^28,29^. It can be quantified by the response time between stimulus and onset of mouth opening or tentacle movement, and by the duration of the mouth-opening time. **F.** Picture of *Hydra* with an opened mouth (white arrow) and tentacles forming a ball shape that is typical of eating behavior. Scale 0.5mm.

Interestingly, microbes present in the glycocalyx are in direct contact with the ectodermal sensory neurons (Fig. 1C)^36^. Previous studies highlighted the capability of *Hydra’s* nervous system to sense and regulate this bacterial community^25,36^. For instance, it was observed that by depriving *Hydra* of its microbial symbionts, spontaneous behaviors such as body contractions become less frequent^37^.

Here, we studied the neuronal activity in freely moving *Hydra* during eating behavior, to uncover the neural circuitry involved. For this, we traced activity of individual neuronal populations using calcium imaging and interrogated their function using cell ablation approach. This revealed that *Hydra*’s eating behavior is controlled by multiple subpopulations of neurons that are activated in a temporally and spatially ordered manner, ultimately leading to mouth opening. In complete absence of microbes (germ-free animals) the mouth opening time is significantly shortened. Adding the full complement of microbes back, restores this defect. Animals’ mono-colonized with the major colonizer *Curvibacter sp.,* also shows severe defect in mouth opening, possibly caused by the production of glutamate. The results demonstrate how, in an animal without a central nervous system, multiple networks of neuronal subpopulations form a neuronal circuit to control a complex behavior. Furthermore, the specific spatiotemporal pattern of neuron activity integrates specific microbial signals, demonstrating that eating behavior does not solely depend on the neuronal state of hunger or satiety: the bacterial community also modulates the neuronal circuits and their state. The evolutionary importance of these observations is discussed.

## Results

### Visualization of the neuronal subpopulations in the head of *Hydra*

First, we confirmed the old observation ^32,33^ that removal of both the body column and the tentacles of *Hydra* polyps did not affect the mouth-opening (see suppl. Video V2), confirming that this property depended on head-specific neuronal regulation.

To visualize the various head specific neuronal subpopulations in *Hydra*, we produced multiple transgenic lines using subpopulation specific genes based on available single cell atlases. For this, we used the promoters of specifically expressed genes, based on available single cell data sets of *Hydra*^24,25^ (See suppl. Fig. S1). The RFamide neuropeptide, (RFa, preprohormone-B, transcript ID^24^: t2059aep) is exclusively expressed in the ectodermal subpopulation N6^38–41^. The genetic target for the ectodermal subpopulation N3 is the neuropeptide Hym-355 (transcript ID: t12874aep)^42^. The marker for the endodermal neuronal subpopulation N4 is annotated as neurogenic differentiation factor 1-like (e-value: 3.03E-140, t14976aep). Their respective promoters were used in expression constructs to either drive the expression of a calcium indicator (GCaMP6s)^26,43^ or for the NTR-MTZ cell specific ablation approach (NTR-MTZ, explained below)^44,45^.

Microscopic investigations of transgenic lines confirmed that the neuronal N6 subpopulation consists of two morphologically different cell types: sensory cells present in a dense cluster in the tip of the head where the mouth will form during feeding, and ganglion cells located in small packages at the basis of the head, close to where the tentacles originate (Fig. 2A-G). These two neuronal N6 types are interconnected by neurites that form radial connections (Fig. 2B). The ganglion cells are frequently circularly connected as well (Fig. 2B, F). A two-dimensional density plot confirmed the concentrated presence of N6 cells at the tip and the base of the head (Fig. 2G). In contrast, N3 neurons are found throughout the body (Fig. 2H). The spatial organization of N3 in the tip of the head is circular around the mouth region (Fig. 2I, K, L). Their density increases at the basis of the head (Fig. 2I-L). N4 neurons in the head reach the highest density at the base and between tentacles, while around the mouth their density is lower (Fig. 2M-S). The morphology of N4 neurons differs between body parts: around the mouth, the N4 neurites form a spider-web structure (Fig. 2R) with a morphology very similar to sensory cells (Fig. 2O, Suppl. Video 6). We call these N4 neurons sensory-like cells.

**Figure 2.**
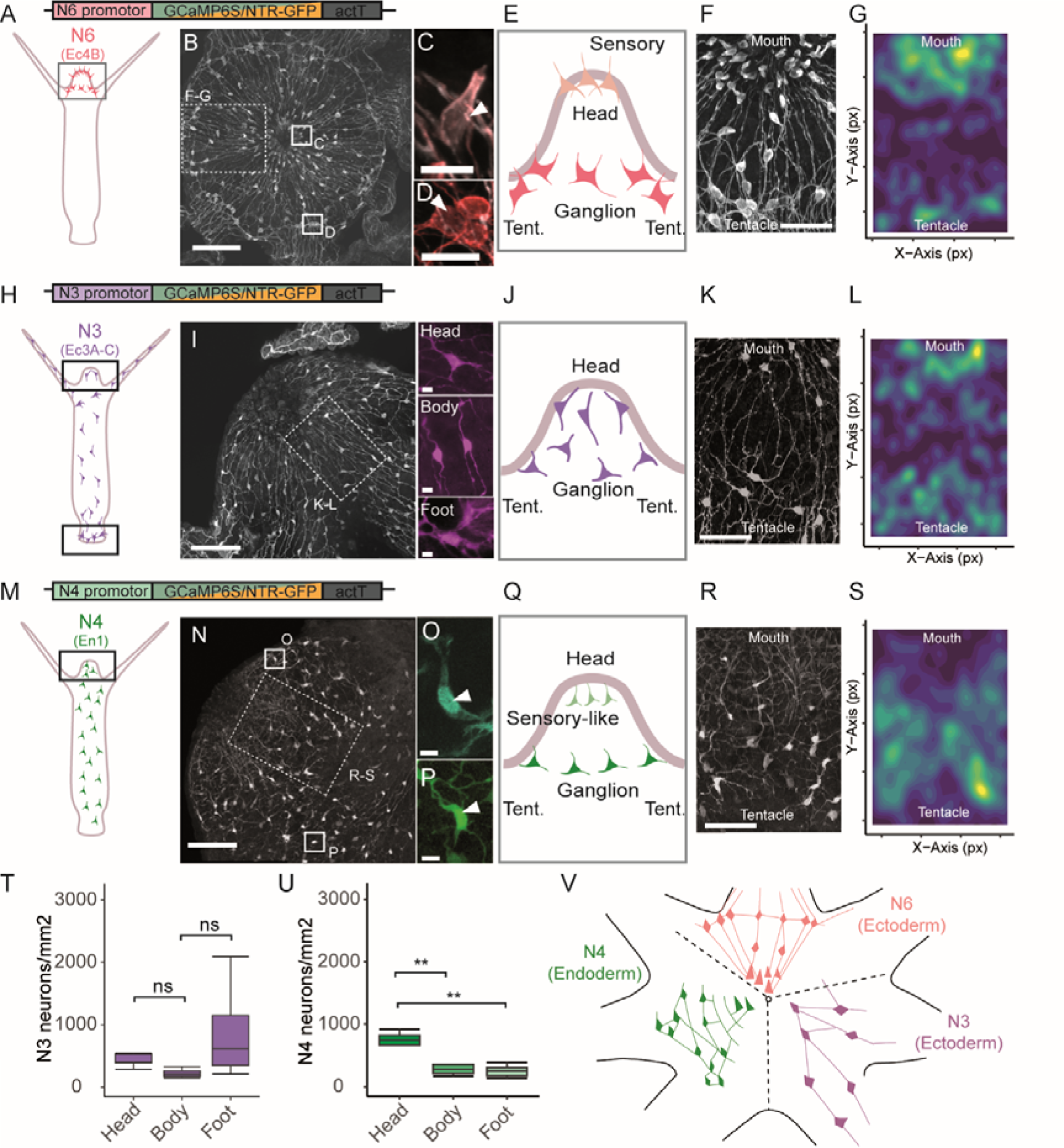
Visualization of the neuronal subpopulations in the head of *Hydra*. **A-G**. Distribution, structure and morphology of ectodermal neuronal subpopulation N6. The constructs used for visualization and manipulation of N6 contains the promoter from an RFa neuropeptide (t2059aep, HVAEP9.T017227.1) regulating the expression of either GCaMP6S or NTR-GFP. **A:** Schematic of *Hydra* and the localization of N6 neurons. **B:** Immunohistochemistry of N6 neurons in the head (scale 100µm) stained with antibodies against GCaMP6S/GFP. **C-D:** Staining (artificial color added) of two representative enlargements showing the two different types of N6 neurons, with sensory neurons at the head tip (**C**) and ganglion neurons (**D**) found in groups around the head base (scale 10µm), as schematically presented in **E**. **F**: enlarged section of **A** with the neurites connecting the neurons (scale 50 µm). **G:** 2D-density plot of the distribution of neurons in a slice of the head (n=5). Higher densities of N6 neurons are present near the mouth and near the basis of the tentacles. **H-L**. Ectodermal neuronal subpopulation N3, for which the construct included the promoter of the neuropeptide Hym-355 (t12874aep, HVAEP2.T004115.1). **H:** Schematic of the localization of N3 neurons in the head (**I**, scale 100µm) and in the body, tentacles and foot (scale 10µm). Their distribution in the head is summarized in **J,** with an enlarged section shown in **K** (scale 50µm). Higher densities are present in the tip and basis of the head (**L**, n=12). **M–S.** Endodermal neuronal subpopulation N4 with the construct containing the promoter of a NEUROD1-like protein (t14976aep, HVAEP4.T008286.1). **M**: the localization of N4 neurons in the polyp. **N:** Overview of N4 neurons in the head (scale 100µm). **O-P:** Staining (artificial color added) of two representative enlargements showing the two different types of N4 neurons, with sensory-like neurons (**O**) and ganglion neurons (**P**, scale 10µm). In the head region, they are mostly present at the basis of the tentacles **(Q, R,** scale 50µm) with a lower density at the tip of the head (**S**, n=10). **T-U.** Density of neurons/mm^2^ in head, body and foot, for subpopulation N3 (**T**) and N4 (**U**). Highest densities of the latter are found in the head (p<0.01, n=5-11, ANOVA, Turkey post-hoc test). **V.** Schematic representation of the distribution and organization of the different neuron subpopulations in the head, from tip (center) to tentacle base. The overlapping locations of the three subpopulations are separated here for clarity. * p≤0.05; ** p≤0.01; *** p≤0.001

The density of N3 neurons was found the highest in the foot, followed by the head, whereas in the body column their density was lower (Fig. 2T). In contrast, most N4 neurons were present in the head with lower and similar numbers in body and foot (Fig. 2U). The distribution of the three neuron subpopulations (N6, N4 and N3) and their neurite networks in the head is summarized in Figure 2V. The individual subpopulations have a clear spatial distribution creating an ordered structure and resulting in a network of nerves with a relatively high complexity in the head region.

### N6, N3 and N4 neurons are differentially active during the mouth opening

Animals were observed during mouth opening as part of their eating behavior and neuronal activity of head neurons was analyzed. The signals for the distinct sensory and ganglion cell types of N6 and N4 were recorded separately. Following the glutathione (GSH) stimulus, the mouth started to open by contraction of the epithelia^32^, which was recorded by plotting mouth width (Fig. 3A-C).

**Figure 3.**
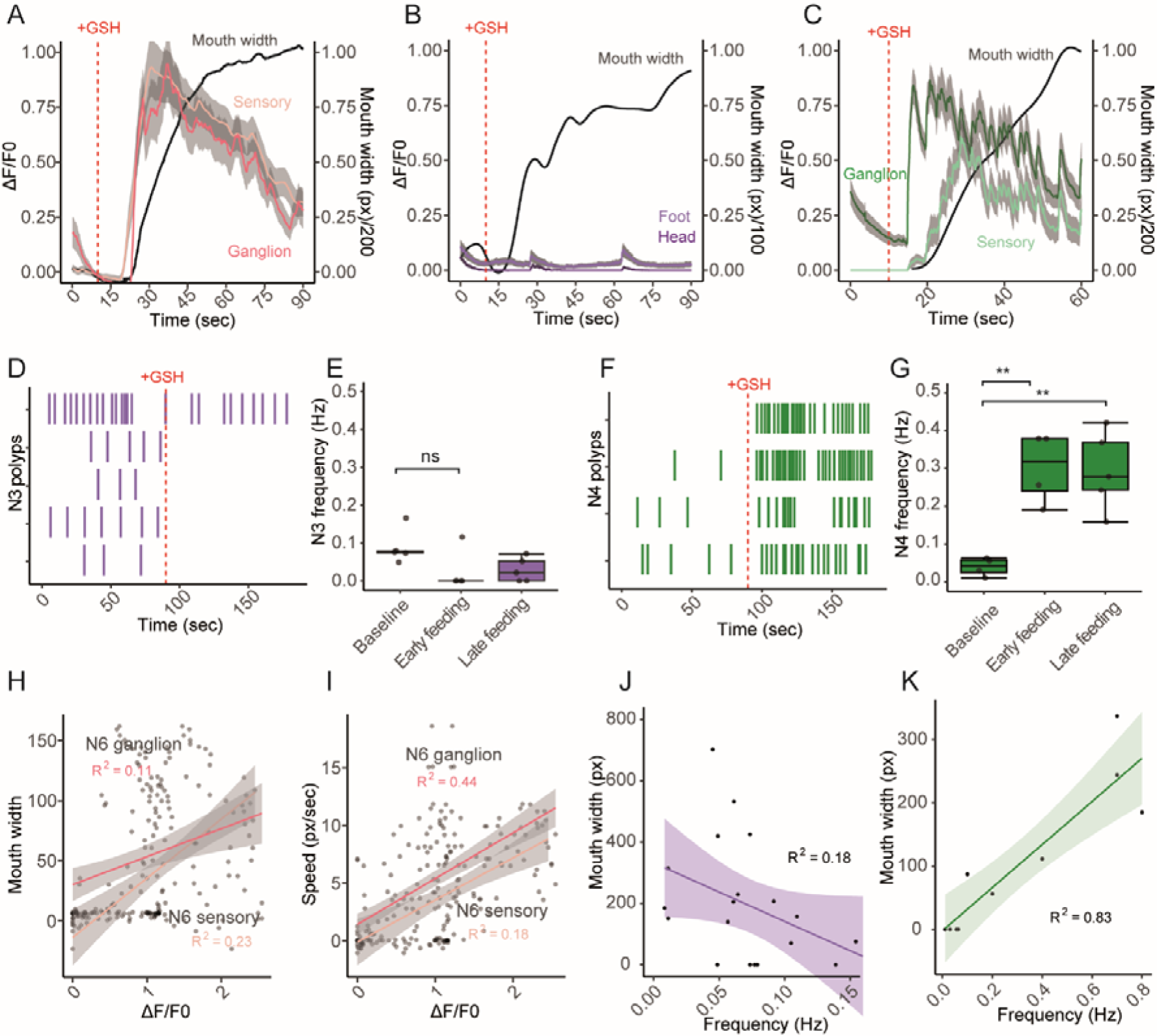
Neuronal response during the eating behavior. A-C. Response of neuron subpopulations to a glutathione food stimulus. **A**: N6 neurons were differentiated into the sensory (red line) and ganglion cells (orange line). The sensory cells responded before the ganglion neurons did. Lines represent the mean of either sensory or ganglion neuronal population from one representative animals with grey shading showing the standard deviation (see suppl Fig. 2 for more animals). At the same time, mouth width was recorded (black line, in pixel). **B: t**he spiking activity of N3 neurons in the head and foot was less obviously affected by glutathione administration. A slightly higher fluorescence change and baseline activity was recorded for neurons in the foot than in the head (one representative animal, see suppl Fig. 3 for more animals). **C:** N4 neurons in the head responded strongly to glutathione administration, with a delay for the sensory cells (mean of population: light green line, light grey shading), whose reaction was also weaker than for the ganglion N4 neurons (dark green line, dark shading, one representative animal, see suppl. Fig. 4 for more animals). **D-G.** Spiking frequency of N3 and N4 neurons before and after glutathione administration. **D**: the spiking activity of N3 in 5 individual polyps decreased in frequency or stopped altogether after the GSH stimulus. **E**: the spiking frequency of N3 neurons at baseline (90s before glutathione, n=5) was lowered during the early feeding response and mouth opening (0-90s post glutathione, n=3) and was restored during the later feeding response (330-420s post glutathione, n=5). **F:** the spiking frequency of N4 neurons increased dramatically after glutathione administration (n= 4). **G**: This increase compared to baseline (n=4) was highly significant during early feeding (n=4, p<0.01, ANOVA, Turkey post-hoc test) and persisted during the late feeding response (n=4, p<0.01, ANOVA, Turkey post-hoc test). **H-K.** Linear correlations of neuron activity onset (bin=30s) with either mouth width or mouth opening speed. A positive correlation was found between N6 ganglion and sensory cells with mouth width (**H**, n=7) as well as with mouth opening speed (**I**). A negative linear correlation was observed between spiking frequency of N3 neurons with mouth width (**J,** n=15). A positive correlation existed between N4 neurons firing and mouth width (**K**, n=6). * p≤0.05; ** p≤0.01; *** p≤0.001

After GSH stimulation, the first signal was recorded within 16.8±26.5s (n=6) for N6 sensory cells, whereas the N6 ganglion cells responded 9.3±5.3s later (Fig. 3A, suppl. Video 3-4), at which time point the mouth started to open. As mouth opening continued, N6 cells activity slowly decreased (Fig. 3A). This was in stark contrast to the activity of N3 neurons, which at first sight seemed unresponsive to the GSH stimulus, both in the head and the foot region (Fig. 3B, suppl. Video 5-6). The N4 neurons responded strongly to the GSH stimulus. A faster response was observed for the N4 ganglion cells located at the base of the head, with a slower and slightly weaker response of the N4 sensory-like neurons (Fig. 3C, suppl. Video 7-8). After the delayed response of the N4 subpopulation, around 40s, the whole cell population started to spike in a synchronous manner (Fig. 3C, suppl. Fig. 5).

Interestingly, N3 neurons responded opposite to N4 to GSH stimulus, as their spiking frequency decreased (Fig. 3D-E). In individual polyps with a relatively frequent N3 spiking at the baseline (Fig. 3D), this became less frequent after glutathione administration. In individual polyps with a low baseline frequency, the firing of N3 neurons stopped completely (Fig. 3D). This was restored to higher frequencies in the late phase of feeding (330-420s post-stimulus). In contrast to N3, the firing frequency of N4 neurons dramatically increased in response to GSH (Fig. 3F) and remained high during late phase of feeding (Fig. 3G). Since N6 cells did not produce pulses but fired more or less continuously (Fig 3A), an analysis of the spiking frequency could not be performed.

Next, we assessed whether there was a correlation between the mouth opening dynamics and neuronal activity. For this, a 30s window starting at the onset of neuronal activity was used and the determined change of fluorescence was correlated with either the mouth width (measured in pixel, px) or speed of mouth opening (px/sec, Fig. H-K). A linear positive correlation was observed for both effects, in sensory as well as ganglion N6 cells (Fig. 3H-I). Fitting the N6 data in a linear correlation for mouth width was better for the sensory cells than for the ganglion cells (R^2^=0.23 and 0.11, respectively, Fig. 3H), however, for mouth opening speed the ganglion neurons fitted better (R^2^=0.44 vs. R^2^ 0.18, Fig. 3I). This indicates that N6 sensory cells were more likely involved in the mouth opening event, while the N6 ganglion cells might be associated with the speed of the tissue movement. The negative correlation between N3 neuron firing and mouth width (Fig. 3J) suggested that a higher frequency of N3 neuron firing correlated with a smaller to no mouth opening. The positive correlation between firing of the synchronous N4 neuron population and mouth width fitted with the highest correlation (R^2^=0.83, n=4, Fig. 3K).

In combination, these data suggest that during eating behavior, N6 sensory neurons are active as the mouth is opening. The activity of N6 ganglion cells correlates with the speed of tissue movement during mouth opening. The spiking of N3 decreases during eating while N4 cells fire more frequently, sending synchronized pulses through the complete polyp.

### Multiple neuronal subpopulations are involved in eating behavior

To identify the contribution of individual neuronal subpopulation in the eating behavior we used the NTR-Mtz cell ablation system. Genetic constructs were used that contained nitroreductase (NTR) fused to GFP, to convert metronidazole (Mtz) to a toxic product that induces apoptosis in the target cell population (suppl. Fig. 6A)^45^. As a control, *H. magnipapillata* strain Sf1 polyps were included that lacks interstitial cells (neurons, nematocytes, gland cells and germline) after application of a heatshock^46^.

The N6 specific promoter caused a strong expression of the NTR-GFP fusion protein in Rfa-positive cells in the polyp’s head (Fig. 4A). Indeed, 93% of Rfa+ cells were also GFP positive, indicating that the N6 line was nearly fully transgenic. Incubation with 10mM Mtz eliminated the N6 neuronal subpopulation within 12h (Fig. 4A, suppl. Fig. 6B). Other neuronal populations remained intact, for instance Rfa+ cells in the tentacles remained detectable, demonstrating that the cell ablation was specific for the target N6 subpopulation. Mtz treatment of control animals containing the GCaMP6S construct had no effect (Suppl. Fig. 6D, G). Despite the absence of N6 neurons, the transgenic animals developed normally (Fig. 4B, compare the polyp pair to the left, without and with Mtz treatment). Similar transgenic animals were produced for ablation of N4 and of N3 (Suppl. Fig. 6D-I). Polyps lacking N4 neurons that are normally present in head, body and foot, developed with an inflated body shape (Fig. 4B, middle pair) and animals lacking N3 neurons (expressed in all body parts) were fully contracted (right-hand pair).

**Figure 4.**
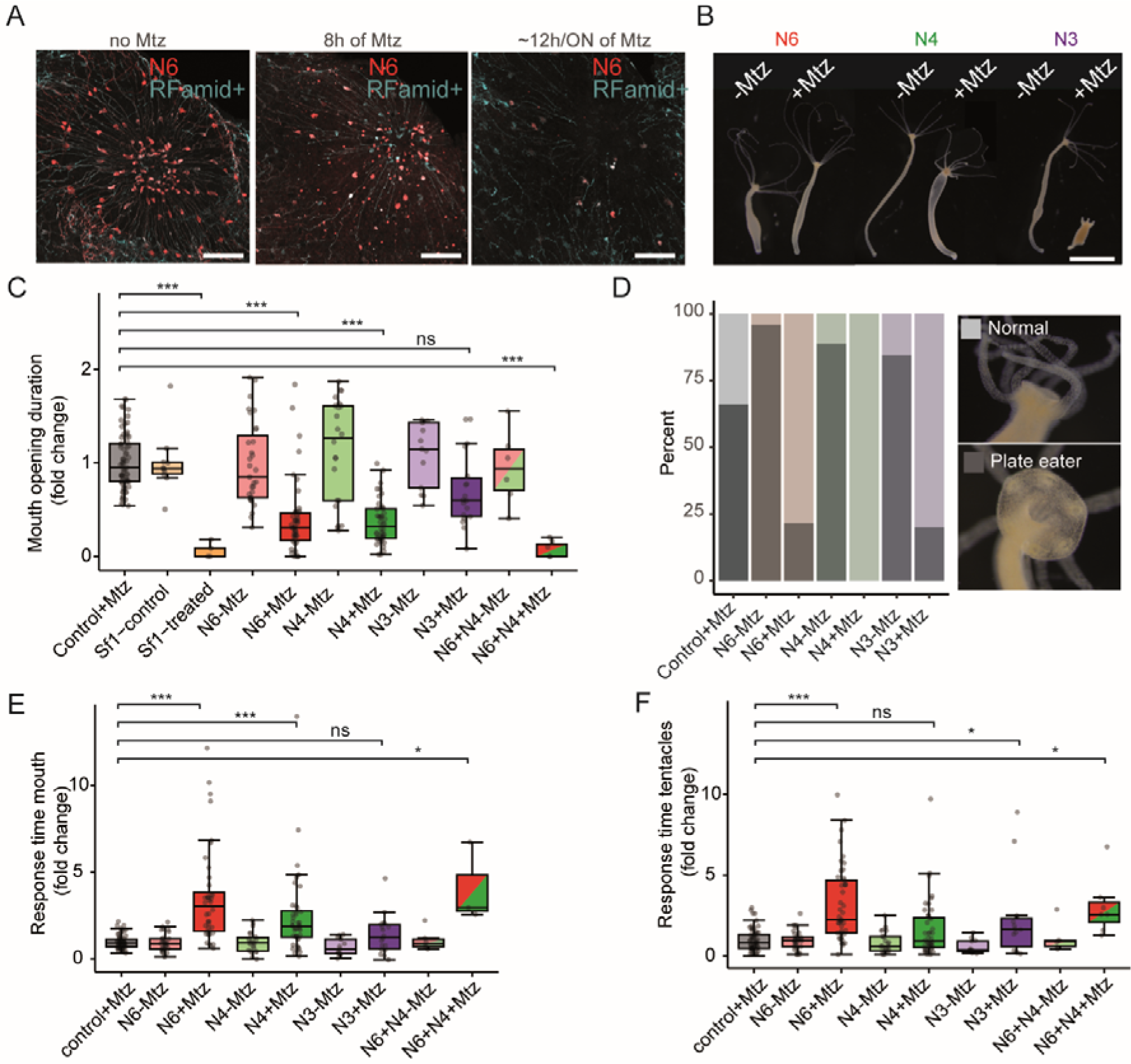
Ablation experiments highlight specific roles of the neuronal subpopulations. **A.** Immunohistochemistry staining for GFP under the promoter for N6 (red) and RFamid (turquoise; expressed in N6 and other neurons) of polyp heads in absence and in presence of 10mM metronidazole (Mtz, 8h and 12h/overnight (ON)). Scale bar 50µm. Note the depletion of N6 neurons over time. **B.** Different NTR transgenic lines of *Hydra* in absence and presence of 10mM Mtz for 12h. Note the inflated body shape following ablation of N4 and the fully contracted body in absence of N3. **C.** The effect of ablating neuronal subpopulations on the mouth opening time. The measured mouth opening time was normalized to the mean response of the control within each experiment before pooling all data. The presence of i-cells is essential for mouth opening. Absence of N6 and N4 significantly (p<0.001, N6: n=46; N4: n=44, N4+N6: n = 7) decreased mouth opening, and when lacking in combination it abolished the behavior. Ablation of N3 had no significant effect (p>0.05). Treatments were compared to the “control+Mtz” group. **D.** The percentage of animals displaying ‘plate eating’ behavior decreased when neuronal subpopulations were ablated (n=18-44). The ablation of N4 inhibited this behavior completely. **E.** The mouth opening response time after administration of GSH was delayed following ablating of neuronal subpopulations. N6 and N4, alone or in combination, had a strong impact on the response time, but N3 did not. (N6: n=46; N4: n=44; N3: n =18; N4+N6: n = 3) **F.** The tentacle movement response time was also affected by ablating the neuronal subpopulations, in particular by N6 (N6: n=46; N4: n=44; N3: n =18; N4+N6: n = 6). All statistical analyses are based on Kruskal-Walli’s rank sum test and Dunn test as post-hoc with Bonferroni method; * p≤0.05; ** p≤0.01; *** p≤0.001.

The effect of apoptotic removal of these different neuronal subpopulations on the eating behavior of the polyps was studied in freely moving *Hydra* individuals (Suppl. Fig. S7, Suppl. Video1). Following GSH stimulation, the duration of the mouth opening period was recorded, as well as the response time required to initiate tentacle or mouth movement. Results were reported as fold-change compared to control (Fig. 4C, E, F). Interestingly when using GSH as artificial food stimulus polyps attempted to ingest the chamber surface (Fig. 4D). This was additionally scored as ‘plate eating’ and described as an extremely wide mouth opening.

As expected, presence of neurons is a pre-requisite for the eating behavior, as their absence in heat-treated Sf1 animals abolished mouth opening completely (Fig. 4C). Ablation of either N6 or N4 subpopulation resulted in a severe reduction in mouth opening time (fold-change compared to control: N4: 0.31±0.398, n=44, p<0.0001; N6: 0.32±0.233, n=46, p<0.0001; Fig. 4C). Removal of N3 caused a non-significant reduction of mouth opening time (0.599±0.94, n=18, p>0.5). When N4 and N6 neurons were removed in combination, the transgenic animals completely stopped opening their mouth (0±0.09, n=7; Fig. 4C).

Plate eating was observed in 65% of control animals when the freely moving polyps spread their mouth wide over the surface of the chamber (Fig.4D). Ablation of N4 neurons completely inhibited this extreme wide opening of the mouth (Fig. 4D).

The response time to open the mouth after the GSH stimulus was affected by removal of N4 and N6 neurons but not by removal of N3 (Fig. 4E). Absence of the subpopulation N6 also strongly delayed tentacle movement (p<0.001, n=46, Fig. 4F).

Taken together, the data show that mouth opening duration, its response time and the response time for tentacle movement during eating behavior are all controlled by two distinct neuronal subpopulations N4 and N6, with a degree of redundancy that adds some resilience to the functioning of this fitness-relevant and important behavior as single ablation could not completely inhibit mouth opening.

### A global neuronal network of connected localized neuron subpopulations regulates epithelial contraction

To investigate if neuronal subpopulations form synaptic-like connections, immunohistochemistry was performed with antibodies targeting the combined RFa+ neuronal subpopulations N1, N6 and N7, or the transgenic lines expressing GFP (Fig. 5). This uncovered that N3 is connected to multiple other ectodermal neuronal subpopulations (Fig 5A-C). Contacts suggestive of synaptic-like structures between N3 and N6^RFa+^ neurites were identified in the head (Fig. 5A), while in the foot N3 and N1^RFa+^ neurons were in close proximity (Fig. 5B). In the tentacles N3 was aligned in nerve bundles with neurites in contact with N7^RFa+^ sensory neurons (Fig. 5C). Contacts between ectodermal and endodermal neuronal subpopulations were also identified, despite their separation by the mesoglea (Fig. 5D-F). For instance, we identified potential contact points between endodermal N4 and ectodermal N6^RFa+^ in the head (Fig. 5D-E). However, in the foot region, no contact points between endoderm N4 neurons and the abundant ectodermal N1^RFa+^ could be identified (Fig. 5F).

**Figure 5.**
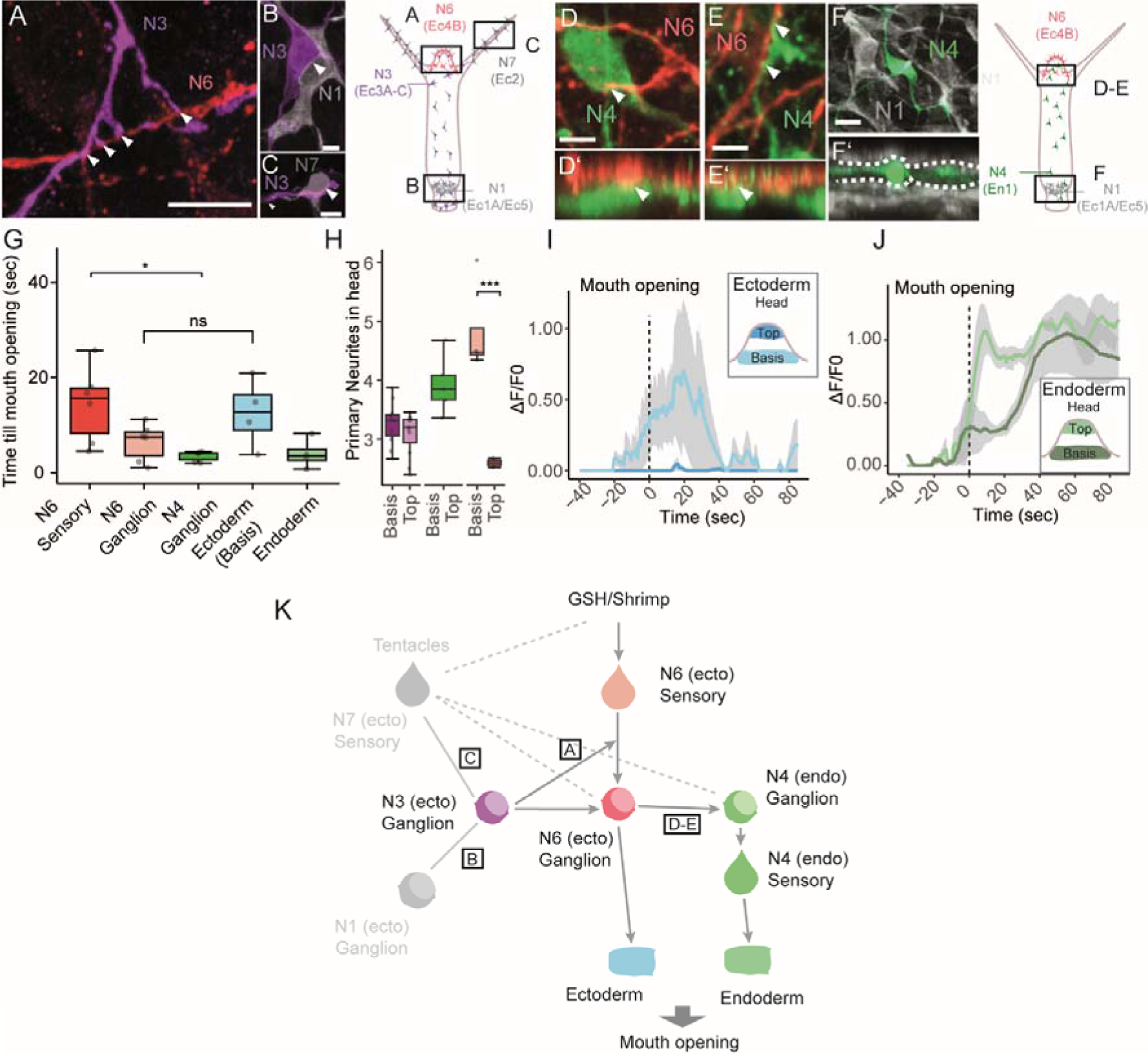
The model of the neuronal circuit controlling the eating behavior in *Hydra*. **A-C.** Identification of contact points between N3 and other ectodermal Rfa^+^ neurons by immunohistochemistry using antibodies targeting N1, N6, N7 in combination, and GFP for visualization of N3. Contact points (white arrows) are present between N3 and N6 in the head (**A**), where N1 and N7 are absent. In the foot (**B**) contact points are found between N3 and N1 and in the tentacles (**C**) they exist between N3 and N7. **D-F.** Potential synaptic contact points (white arrows) were also identified between endodermal N4 and ectodermal neuronal populations at the base of the head. Four examples are shown in **D-E**, where potential contacts between endodermal N4 and N6 (white arrows) were observed. Two zoomed pictures are shown, one frontal maximum projection (D’) and one orthogonal maximum projection (E’, scale bar 5µm). In the foot (F-F’) there was no contact between N4 and N1. **G.** Time gap between the first neuronal activity and the beginning of mouth opening. The N6 subpopulation is split into sensory and ganglion cells. A higher value indicates a faster response to the food stimulus, as seen for N6 sensory cells (n=4-7, Kruskal-Wallis and Dunn post-hoc). **H.** The number of primary neurites of each subpopulation divided into head top and head lower part (basis) for N3 (purple) and N6 (red, orange). The highest number of primary neurites are found for N6 at the basis of the head (n= 4-11, ANOVA, Turkey post-hoc). **I-J.** Contraction response of the epithelia to the food stimulus, with ectoderm (**L**) and endoderm (**M**). The time point when the mouth opened is indicated. No contraction of ectoderm in the head top was identified butt a time relapse between contraction of endoderm at the head top and base is visible **H**. (n=4) **K.** The model of the neuronal circuit involved in the eating behavior. N6 sensory cells detect glutathione first and propagate the signal to N6 ganglion cells, where it spreads to the endodermal N4 ganglion cells. At the same time, the signal propagates to N3 cells which modulate the response and stops firing, leading to mouth opening. Contact between N3 ganglion cells and N1 and N7 neurons ensures further spread of the signal through the body of the polyp. * p≤0.05; ** p≤0.01; *** p≤0.001

To investigate the sequential activity of the neurons in the neuronal circuit, we measured the time gap between the first neuronal activity and the onset of mouth opening. A longer time-gap relates to an earlier response in the eating behavior. Figure 5G shows that the earliest responses were observed for sensory N6 cells, followed by N6 ganglion cells and then N4 ganglion cells (Fig. 5G, n=4-7). This suggests that sensory N6 cells detect the food stimulus first, to pass the signal on to ganglion N6 cells, before the N4 cells respond.

The number of primary neurites located in top part of the head and at its base was determined for N6 and N3 (Fig. 5H). The top of the head contained the fewest N6 neurites, and the base contained the most (Fig. 5H). This would enable a signal picked up by N6 sensory cells to be not only propagated but also enhanced via N6 neurites at the base, where the contact between N6 and N4 cells (cf. Fig. 5H, D-E) ensures involvement of the latter. At the same time, contact between N6 and N3 would allow the inactivation of the N3 cells.

As mouth opening requires the contraction of epithelia, we also measured the time required to initiate contraction of both ectoderm and endoderm involved in mouth opening (see Methods for the application of calcium imaging constructs under control of an actin promoter for this)^47^. First, we observed that the ectoderm of the head base contracted before the endoderm did (Fig. 5I, J). While the endoderm was activated in the whole head region at some point during the behavior, with a faster response at the top than at the base of the head (Fig. 5J), the ectoderm was only active at the base of the head, close to the tentacles (Fig. 5I). The time required for ectodermal contraction at the head base and for endodermal contraction till mouth opening differs.

All data taken together suggest that the reaction flow went from the N6 sensory cells to the ectodermal epithelium and to N6 ganglion cells, and from there to the N4 ganglion neurons and then to the endodermal epithelium. This is summarized in Figure 5K.

### The role of bacteria: mono-association of *Curvibacter sp.* reduces mouth opening

Since there are symbiotic bacteria in the immediate proximity of the head neurons^36^, we next asked whether these bacteria might have an influence on the neuronal circuit identified here that control eating behavior. For this, germ-free (GF) animals were compared with wildtype (Wt) and recolonized with a number of pure cultures of native bacteria as described previously^48,49^. Intriguingly, germ-free animals kept their mouths open much shorter than control animals did (p<0.01, Fig. 6A). Mono-association of polyps with single members of the core bacterial community, (including *Duganella, Pelomonas* or *Undibacterium* species) rescued this defect (Fig. 6A), although monoassociation with *Pseudomonas* or *Acidovorax* had no effect (Fig. 6A). Completely unexpected results were obtained with animals that were mono-associated with *Curvibacter sp.,* which is the most abundant representative in the wildtype *Hydra* AEP microbiota^48–50^. Exclusive presence of these bacteria reduced the mouth opening time to nearly zero (n=47, Fig. 6A-B). The effect could be restored to some degree by co-addition of a second bacterial species, whereby all tested di-associations produced similar effects (Fig. 6B). The combination of *Curvibacter* with *Undibacterium* and *Duganella* restored the mouth opening time to normal (Fig. 6B).

**Figure 6.**
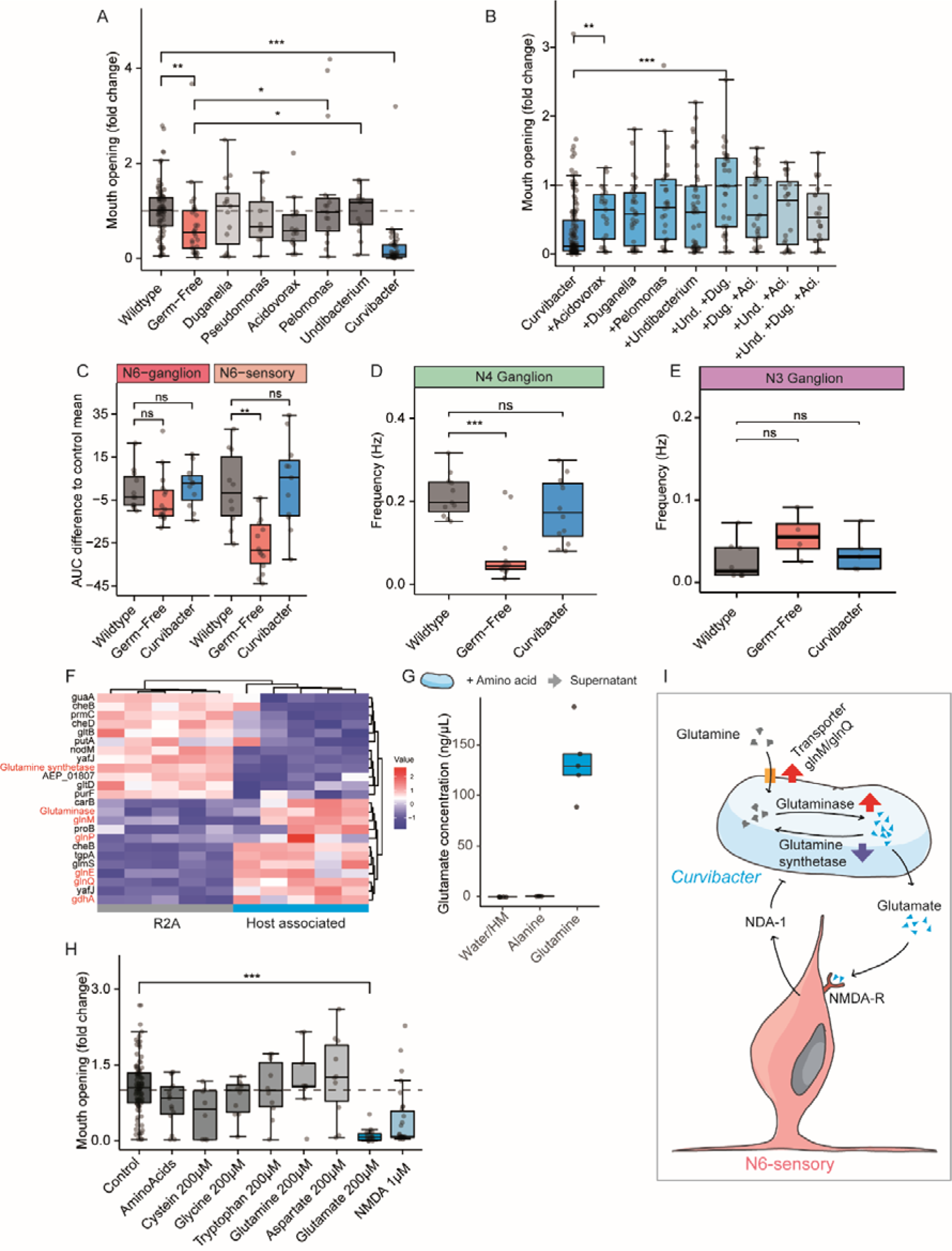
Mono-association of *Curvibacter sp.* inhibits mouth opening in *Hydra* via glutamate production. **A.-B.** Wildtype (Wt) *Hydra* was made germ-free (GF) with antibiotic (AB) treatment and then recolonized with single constituents of its native microbiota or combinations thereof. **A.** Absence of bacteria decreased the mouth opening duration during feeding. This could be partially restored by recolonization with various bacterial species, but mono-association with *Curvibacter sp.* strongly inhibited mouth opening. Results of replicas (n>5) were normalized to the respective control (dotted line: mean of the control) before pooling experiments. **B.** The negative effect on the mouth opening time of mono-associated *Curvibacter sp.* was rescued by other bacterial species with increasing community complexity. Und: *Undibacterium sp*., Dug: *Dugonelia sp*., Ac: *Acidovorax sp*. *Curvibacter sp*. was present in all. **C-E.** Neuronal activity during eating behavior in transgenic animals with wildtype microbiota, germ-free animals or recolonization with *Curvibacter*. **C**: the activity of N6 sensory cells was negatively affected by absence of bacteria, while recolonization of *Curvibacter* rescued this effect. N6 ganglion cells were not affected, Area under the curve (AUC) was calculated by taking the mean of respective controls and subtract this from treatment (n=8-10). **D:** The spiking frequency of N4 was severely impaired in GF polyps which was reversed by presence of *Curvibacter*. (n=10-11)**. E:** Absence of bacteria or mono-association with *Curvibacter* did not affect N3 neural spiking frequency during feeding (one experiment, n= 3-5). **F.** Differentially expressed genes in *Curvibacter sp.* grown in minimal growth medium (R2A) and in mono-association with its host. Multiple genes are involved in glutamine/glutamate binding, transport and metabolism shown in red font. **G.** *Curvibacter sp.* grown *in vitro* in *Hydra* culture medium supplemented with 200µM glutamine secreted glutamate into the medium (n= 5), whereas addition of alanine to the medium had no effect **H.** Adding glutamate to the medium strongly inhibited mouth opening, but other amino acids had no effect. Addition of NMDA mimicked the glutamate inhibition (n=10). **I.** Model of the mechanism how glutamate production by *Curvibacter sp.* affects its host. Red arrows indicate upregulation of bacterial genes during host association, with purple arrows showing down regulation, which in combination lead to higher glutamate secretion. This binds to an NMDA-receptor on N6 sensory cells to produce NDA-1, an antimicrobial peptide that limits *Curvibacter* propagation^36^. Kruskal-Wallis, Wilcoxon test, * p≤0.05; ** p≤0.01; *** p≤0.001

The strong inhibitory effect on the mouth opening time by mono-association of *Curvibacter* led us to investigate the effect of these bacteria on the neuronal activity during eating behavior. For this, the neuronal activity of Wt, GF and polyps mono-associated with *Curvibacter* was compared by calcium imaging (Fig 6 C-E). As shown in Figure 6C, the activity of N6 sensory neurons in germ-free animals was much lower compared to controls. Interestingly this could be restored by presence of the *Curvibacter* symbiont (Fig. 6C). Likewise, *Curvibacter* restored the decreased N4 activity of GF polyps (Fig. 6D). Activity of N3 neurons was not affected (Fig. 6E).

### *Curvibacter sp.* affects neuronal cells by means of glutamate

To investigate the mechanism how *Curvibacter sp*. influences the eating behavior and neuron activity on a molecular level, gene expression pattern of *Curvibacter* in mono-association with *Hydra* was compared to the expression profile of the bacteria cultured in minimal growth medium R2A. Differentially expressed genes were identified and the metabolic pathways in which these genes were involved were examined. This approach identified pathways for alanine, aspartate and glutamate metabolism, among others (Fig. 6F). In particular, genes associated with glutamate metabolism were differentially expressed (Fig. 6F). While glutamine synthetase was downregulated in presence of the *Hydra* host, glutaminase, glutamate dehydrogenase (*gdhA*) and *glnM*, *glnP* and *glnQ* (glutamine transporters) were over-expressed (Fig. 6F). This points to a glutamate production during host-association. Since glutamine binding and uptake associated genes are also upregulated in host-association, we assume that host-associated *Curvibacter sp.* secretes glutamate while taking up glutamine.

Previous work had described a putative NMDA-like glutamate receptor in *Hydra* tissues to be involved in the eating behavior^51–53^. Since N6 sensory cells express the NMDA receptor (Suppl. Fig. 8) we hypothesized that bacterial glutamate might bind to the NMDA-like receptor present on N6 sensory neurons. To test this, *Curvibacter sp.* was first cultivated i*n vitro* in *Hydra* culture medium supplemented with 200µM glutamine. The glutamate concentration in this supernatant reached 129±36.1ng/µL. When *Curvibacter* was cultivated in *Hydra* culture medium with alanine, little glutamate was secreted (Fig. 6G). Next, we tested the effect of glutamate on the eating behavior. As shown in Figure 6H addition of glutamate inhibited mouth opening duration. Addition of NMDA had a similar inhibitory effect, corroborating the hypothesis of an NMDA-receptor being involved. Adding various other amino acids had little to none effect on mouth opening duration (Fig. 6). Taken together and in accordance with previous biochemical and functional evidence of the occurrence of putative NMDA-like glutamate receptors in *Hydra* tissues and with the fact that N6 sensory cells are equipped with this receptor, we assume that *Curvibacter* produced glutamate can affect the N6 neurons via this receptor.

## Discussion

Cnidarians emerge as informative models for neuroscience, as they have surprisingly complex neuronal circuits and enable the study of neuronal activity in complete organisms lacking a centralized nervous system^20,26,54,55^. Neuronal control of behavior in cnidarians is dispersed and control takes place in neuronal subpopulations. Recent work in *Hydra* have highlighted that simple spontaneous behaviors are controlled by single neuronal subpopulations^26^. *Hydra* has multiple non-overlapping neuronal networks, each of which can regulate a single behavior, but they are also collectively involved in mechanosensory processing^56^. *Hydra* is also a well-described metaorganism which is colonized by a stable and functionally relevant microbiota^37,48,50,57–59^. Here we studied the interplay and coordination between multiple neuronal subpopulations and epithelial cells that together with the microbial colonizers are involved in the eating behavior. Our work emphasizes the importance of the microbiota on neuronal circuits. The results offer an opportunity to unravel the evolution of the interplay between bacteria and the nervous system mechanistically.

### The eating behavior requires coordination between multiple neuronal subpopulations

The eating behavior illustrates beautifully how, in the absence of any form of centralization, different neuronal subpopulations interact with each other to coordinate a complex behavior. In the presence of a chemical signal for food, for which we used reduced glutathione, different neuronal subpopulations were either activated or inactivated, to coordinate the epithelial movement leading to mouth opening (Fig. 3, Fig. 5K). First N6 sensory cells at the tip of the head are activated, followed by the N6 ganglion cells at the head base, while the local ectoderm contracts. When N6 ganglion cells are activated, the frequency of N4 spiking is increased and the signal spreads from the base back to the tip of the head, leading to contraction of the endoderm (Fig. 3, Fig. 5K). At the same time, N3 decreases in spiking frequency and the mouth is opening, suggesting these neurons have an inhibitory function on mouth opening (Fig. 3). Ablating N3 neurons did not inhibit the mouth opening, whereas ablating N4 and N6 did (Fig. 4C). Interestingly, these neuronal subpopulations control different aspects of the mouth opening and eating behavior: N4 regulates the mouth opening width (Fig. 4D), whereas N6 is involved in the recruitment of the tentacles (Fig. 4F). In combination, these three different neuronal subpopulations form the neuronal circuit controlling the epithelia involved in the eating behavior (Fig. 5K).

The fact that several non-overlapping neuronal subpopulations are involved to control different aspects of eating behavior suggests that they must be in contact with each other. One of the highest densities of neurons in *Hydra* is at the tentacle-head junctions at the base of the head^60,61^. At this location, the densities of N3, N4 and N6 populations are particularly high, with increased numbers of primary neurites compared to the body column (Fig. 2G, L, S, Fig. 5H). In addition, in this region distinct contact points between neurons of different neuronal subpopulations could be identified (Fig. 5A-F). In combination with the sequence of activity after glutathione stimulus (Fig. 3) this led us to the conclusion that the base of the head is the region where the different signals are being integrated and distributed to the endodermal and ectodermal networks. The complexity of neurites and synaptic structures in this region was already identified by electron microscopy^60–65^. Our results highlight how a relatively simple neuronal network in *Hydra* can result in a stunning complexity in order to process sensory information into multiple responses to control the complex eating behavior. We note potential similarities to neuronal control mechanisms in the jellyfish *Clytia hemisphaerica*, where an apparently diffuse network of neurons is functionally subdivided into a series of spatially localized subassemblies whose synchronous activation controls food transfer from the tentacles to the mouth^54^. However, that organism depends on functional modules, whereas in *Hydra* multiple subpopulations within a single circuit coordinate the behavior.

### Eating behavior becomes severely impaired when the microbiota is disturbed

Polyps mono-colonized with *Curvibacter sp.* had drastically reduced mouth opening time, suggesting a bacterial signal interfered with the neuronal circuit that controls this fitness relevant behavior. This striking inhibitory effect was all the more surprising, as *Curvibacter sp.* normally represents around 70% of the *Hydra* bacterial microbiota and has not been associated previously with any negative effect on the host^57^. The inhibitory effect of *Curvibacter sp*. could be reversed by increasing bacterial diversity while adding back specific members of the core microbial community (Fig. 6B).

The inhibitory effect of *Curvibacter sp*. on eating behavior was not accompanied by a detectable change in neuronal activity compared with the control (Fig. 6C-E). Instead, mono-association of *Curvibacter sp.* reversed the effect of germ-freeness back to control conditions. This highlights that *Curvibacter sp.* affects neuronal activity, and also that neurons are able to sense the presence of *Curvibacter* presence. Since the N6 sensory neurons are in close contact with the microbiota^36^, their response was to be expected, but a similar effect on the endodermal N4 population (Fig. 6D) was rather unexpected and suggest that *Curvibacter sp.* has a more global effect on the nervous system.

The transcriptional response of *Curvibacter sp.* to the host environment points to the secretion of glutamate in the presence of glutamine, which was supported by *in vitro* observations (Fig. 6G-I). The neuronal subpopulations N3, N4 and N6 express an NMDA receptor that could responds to bacterial glutamate and integrates this information into the neuronal circuit of the eating behavior (NMDAR and mGlu, see Suppl. Fig. 8). Since N4 and N6 but not N3 neurons showed a response to *Curvibacter sp.*, we assume that N4 and N6 receive and integrate the bacterial signal. Our work shows that the old observation published by Lenhoff (1961)^52^ that glutamate has a negative effect on eating behavior in *Hydra* may find its explanation in the microbial colonization of *Hydra*.

### Evolutionary perspective

Altogether, our findings confirm and expand on the idea that in animals without a central nervous system, a complex behavior is controlled by multiple subpopulations of neurons, forming circuits and modules^54^. Our observations presented here show that this not only requires the coordination of multiple neuronal circuits, but also that signals from the microbial environment play an important role. We present data that support a model (Fig. 6I) in which in the critical phase of mouth opening, can be affected by microbially produced glutamate.

Already in 1963, the evolutionary biologist Tinbergen outlined an organizational framework that would control complex behavior^66,67^. His research involved four levels of analysis: phylogenic, developmental, functional, and mechanistic investigations. Our findings of the influence of the microbiota on the neuronal control of *Hydra*’s eating behavior, which co-evolved with this host^49,57^, adds this as an additional environmental perspective to be considered when studying complex behavior.

That bacteria are able to produce molecules that are active on neuronal cells has been known for quite some time^68,69^, but most work has been carried out in mammals. Here we show that the integration of bacterial signals into neuronal circuits might be as evolutionary ancient as the first nervous system, as it already exists in cnidarians. Our observation that bacterial glutamate plays a crucial role in this interaction, together with the numerous findings on the influence of this molecule on mammalian intestinal physiology^9,69^, support the idea that it is part of an ancestral interkingdom language.

## Acknowledgements

This work was supported in part by grants from the Deutsche Forschungsgemeinschaft (DFG), the CRC 1182 “Origin and Function of Metaorganisms” (to TCGB.) and the CRC 1461 “Neurotronics: Bio-Inspired Information Pathways” (Project-ID 434434223 – SFB 1461) (to TCGB and AK). T.C.G.B. appreciates support from the Canadian Institute for Advanced Research. AK is supported by a DFG grant KL3475/2-1. C.S. and T.S. acknowledge funding by the DFG under Germany’s Excellence Strategy 2082/1-390761711 (3D Matter Made to Order). We thank Trudy Wassenaar for critical reading of the manuscript. We thank the members of the Bosch lab for support and discussion, and Andreas Tholey, Christoph Kaleta, Georgios Marinos and Karlis Moors for discussion. We also thank Urska Repnik and Marc Bramkamp from the Central Microscopy Facility at the Biology Department of the University of Kiel for excellent technical support. We highly appreciated the expertise provided by the sequencing facility at the Institute of Clinical Molecular Biology (IKMB) in Kiel, Germany.

## Authors contribution

C.G. and T.C.G.B. conceptualized the project and wrote the manuscript. T.C.G.B., J.W., A.K., Y.G., D.P. and C.G. designed and performed experiments on transgenesis. T.C.G.B., D.P., C.S., T.S., E.H. and C.G. designed and performed histological, behavioral experiments. C.G., E.H., T.L. and T.C.G.B. designed and performed neuronal activity and microbiota experiments. T.L. and C.G. analyzed the data.

## Data and code availability

• Source data reported in this paper will be shared by the lead contact upon request.

• Codes used for the analysis and statistical analysis will be shared by the lead contact upon request.

• Any additional information required to reanalyze the data in this paper will be shared by the lead contact upon request.

## STAR Methods

### Materials availability

The plasmids and transgenic *Hydra vulgaris* AEP generated in this study are available upon request.

### Code availability

All codes used in this study are available upon request.

## Experimental Procedures

### *Hydra* maintenance

In this study used *Hydra* polyps (*Hydra vugaris* AEP, *Hydra magnipapillata sf1*) were cultured according to standard procedures in standard *Hydra* culture medium (CaCl_2_ 0.042g/L; MgSO4x7H_2_0 0.081g/L; NaHCO_3_ 0.042g/L, K_2_CO_3_ 0.011g/L in dH_2_O) ^70^. The animals were kept in 250mL glass beaker at 18°C with a 12/12h light cycle. The feeding regime was strictly three times per week with *Artemia* nauplii for at least two weeks before any experiment. Animals were starved for 1-3 days before either an ablation experiment or a calcium imaging analysis. There was no difference in the mouth opening duration between 1-3 days of starvation.

### Generating germ-free animals and re-colonization

Germ-free animals were derived by treating animals for five days with an antibiotic cocktail containing rifampicin, ampicillin, streptomycin and neomycin in final concentrations of 50 µg/ml each and spectinomycin of 60 µg/ml, as previously described^49^. Control polyps were incubated in 0.1% DMSO for the same time since rifampicin is dissolved in DMSO. The antibiotic cocktail was replaced after 72h of incubation. After 5 days in antibiotics, the animals were transferred to sterile *Hydra* culture medium and incubated for another 2 days. On the second day in sterile *Hydra* culture medium, animals were recolonized with defined bacteria or communities and medium was exchanged. After another 3 days of incubation with defined bacteria or communities, polyps were used for the behavioral assays or RNA sequencing. The germ-free status was checked twice during the protocol, on the seventh day and the tenth day of the protocol via plating macerated polyps on R2A-agar plates. Random samples were also tested via PCR using universal rRNA primer Eub-27F and Eub1492R^71^. No colonies formed on the R2A agar plates after one week of incubation at room temperature and absence of amplification product confirmed the germ-free status.

Germ-free animals were monocolonized with pure bacteria cultures of the core members of *Hydras* microbiota: *Curvibacter* AEP 1.3 (NCBI:txid1844971), *Duganella* C 1.2 (NCBI:txid1531299), *Undibacterium* C 1.1 (NCBI:txid1531302), *Acidovorax sp.* AEP 1.4, *Pelomonas* AEP 2.2 (NCBI:txid1531300) and *Pseudomonas sp.*^50^. Bacteria were cultured from existing isolate stocks in R2A medium at 18°C for three days and subcultured the day before recolonization (dilution depending on the bacterium). In all experiments we started from a fresh cryostock and identity of bacteria was regularly tested. From the overnight culture approximately 10^5^-10^6^ cells were added to the 50mL sterile *Hydra* culture medium containing 30-50 animals. For the different combinations of bacteria, each bacterium was added in at equal ratios. After three days the recolonization success was checked by plating three macerated polyps per treatment in a 1:1000 dilution on R2A agar plates and counting the CFUs after three to four days of incubation at 18°C. Recolonized animals were only used when recolonization was successful and in agreement with previous published values^72^.

### Promoter identification and extraction

Marker genes specifically expressed in the different neuronal subpopulations were identified using the single cell atlas previously published ^24,25^ (suppl. Fig. 1). Genes were then mapped against the different available genomes of *Hydra* (nih.gov/HydraAEP) and their promotor were extracted as 1000-1500bp upstream of the gene, by including the first 30bp of the open reading frame. The sequence was than cloned into pGem-T Easy (Promega, cat# A1360) while restriction enzyme binding sites were inserted to further clone the construct into the LigAF vector (for sequences see suppl. Table 1).

### Transgenesis and constructs

Transgenic *Hydra vulgaris* AEP were derived following the established protocol by Wittlieb *et al*.^70,73^ using a modified version of the LigAF vector. Different lines were produced in which the specific promotors for desired expression in the neuronal subpopulation regulated either GCaMP6S (as in Dupre *et al*. ^26^) with an actin terminator or the nitroreductase (NTR)^44^ (*in silico* codon optimized) coupled to an eGFP at the C-terminus followed by an actin terminator sequence (see suppl. Table 1 for sequences). As previously described, the construct was injected via microinjection in embryos resulting in mosaic animals. Animals were screened for transgenic neurons and selected to produce fully stable transgenic animals. We then induced embryogenesis in the transgenic lines and derived F1-generations which ensured that the construct was incorporated in all cells. This was successful for transgenic lines N4 and N6 while for N3 reached a non-mosaic stable population only (see suppl Table 1).

### Histology

For antibody staining, *Hydra* polyps were relaxed with 2% urethan(Sigma-Aldrich, U2500) in *Hydra* culture medium for less than 2min and fixed for 2h (RT) or overnight (4°C) in Zamboni (Morphisto, cat#12773). Following 3 washes in PBS with 0.1% tween (PBST) followed by an incubation in PBS with 0.5% TritonX100 and an 1h of blocking in PBST with 1% bovine serum albumin (BSA, Roth, cat# 8076.1). Animals were than incubated overnight at 4°C with the primary antibody in PBST and 1% BSA. Primary antibodies used in this study were: anti-GFP (Biozol, cat# GFP-1010, 1:1000 dilution) and anti-FMRFamid (BMA Biomedicals, cat# T-4322, 1:1000 dilution). After the primary antibody incubation, four 15min washes in PBST with 1%BSA were performed before adding the secondary antibody. Secondary antibodies used in this study were: goat anti-chicken Alexa Fluor 488 (Invitrogen, cat# A11039, 1:1000 dilution) and donkey anti-rabbit Alexa Fluor 546 (Invitrogen, cat# A10040, dilution 1:1000). Animals were incubated for 2h at RT with the secondary antibody. After the secondary antibody another four 15min washes in PBST (here 0.5% tween) with 1% BSA were performed followed by a short 5min incubation in TO-PRO™-3 Iodide (642/661)(Invitrogen, cat# T3605, 1:1000 dilution). The animals were mounted in moviol with DAPCO on glass slides and stored at 4°C till imaging.

### Imaging and analysis

Fixed and stained animals were imaged either with a LSM900 (Zeiss) or Axio Vert.A1 (Zeiss) using colibri 7 (Zeiss) as a light source. Further processing of the images was performed with Zen Blue 3.4 software (Zeiss) or Fiji^74^. For the analysis of neuronal densities and distribution we used the Cell Counter plugin by Fiji. For counting, a rectangular area was subsampled from the images to count comparable areas (see Fig2B, I and N). For the densities, we calculated the density of neurons per mm^2^. For the 2D density plots (Fig 2G, L and S) we aligned the rectangle area using Fiji and extracted the x- and y-coordinates. Data were analyzed using R (v4.0.3)^75^ over RStudio IDE^76^ and for the visualization the plugin tidyverse (v1.3.1)^77^ was used. For the characterization of the primary neurites, we counted all neurites originating from a neuron soma. In all cases at least five animals were analyzed. Multicolor images shown throughout are pseudo-colored composites (maximum projection), with brightness and contrast adjusted for clarity.

### NTR and sf1 cell ablation experiments

Animals were incubated overnight in 10mM Metronidazole (Sigma, cat# M1547)^44,45,54^. On the next morning animals were screened under a fluorescence microscope for absence of GFP^+^ cells. Once it was determined that the ablation had been successful, the animals were washed once in *Hydra* culture medium and used for behavioral assays or histology on the same day. Each experiment included a control of *Hydra vulgaris* wildtype and the corresponding GCaMP6S transgenic line with Metronidazole and the NTR-GFP transgenic line without Metronidazole. In all experiments at least 5 animals per treatment were used.

*Hydra magnipapillata* Sf1 were exposed to 28°C for 48h together with a control (*H. magnipapillata*) for the heat shock and afterwards kept for 19 days under standard culture conditions. Neurons were quantified on day 5, 8, 11, 14 and 19 using a cell maceration protocol^22^. Polyps were dissociated in maceration solution (1:1:13, Glycerol, Acidic acid, *Hydra* culture medium) at 32°C for 30 minutes. Afterwards cells were fixed in 8% PFA and spread out on gelatin-coated slides. Counting was done blinded.

## Behavioral analysis

### Acquisition

To analyze the effect of cell ablation and bacteria on the eating behavior, we developed a recording system where we can observe multiple animals at once and animals were minimally restrained. For this, a chamber was used where 5-6 animals could be observed under controlled fluid flow (Suppl. Fig. 7). The chamber consists of a two-piece aluminum case and two plexiglass pieces in which one cavity was milled and the other used as a lid (see suppl. Fig. 7). These were connected and liquid tight via braces. Animals could survive in the chamber for weeks as long as fresh *Hydra* medium was supplied. The chamber has a height of 0.4mm and two channels on both sides fitted with tubes through which medium can be manually supplied. The animals were recorded at 18°C in an insulated climate chamber to avoid external stimuli using M3C Wild Heerbrugg binocular microscopes and Axiocam 208 color (Zeiss), taking a picture every 2 seconds.

### Mouth opening, tentacle response and analysis

The animals were given 10 min to adapt to the recording chamber before recording started and another 5-10 min before reduced glutathione (GSH, Roth, cat#6382.1) was supplied via the tube system. In all assays a final concentration of 10µM GSH was used, prepared in the same medium as the animals were kept in prior to the experiment using a 0.1M stock solution. Each animal was only recorded once. Acquired movies were blinded to their treatment and assigned with a random three-digit number before analysis. The behavior was manually annotated. For the following different behaviors: the mouth opening time, tentacle movement and the type of mouth opening (see suppl Video 1). As some animals exhibited multiple mouth openings during the assay, for the mouth opening time only the first event was recorded. The raw data from the video analysis were further normalized by the mean of the respective controls within each experiment to obtain the fold-change information between treatments. The data were merged for analysis and plotting.

### Mouth opening width

In order to correlate the mouth opening behavior and neuronal activity, we measured the width of the mouth opening during GCaMP6S recording via automated tracking of the opposite edges of the mouth. This tracking was done using icy^78^ and the tracking plugin^79^. Afterwards tracks were manually cleaned, and missing links were integrated. Using the track manager with the integrated function “Distance profiler” the distances between the two different tracks were calculated in pixel. The tracks were then smoothed using the integrated ksmooth function in R^75^. As the mouth opening onset to analyze the time sequence of neuronal activation before mouth opening, the first increase in the slope was taken after the addition of GSH and where there is no decrease within a 20 sec window.

### GCaMP6S imaging acquisition and calcium traces extraction

To analyze the neuronal activity during the eating behavior, we developed a system to record freely moving animals while adding GSH. The animals were placed in commercially available channel slides with a height of 0.2mm and a width of 5mm (Suppl. Fig. 7.; Ibidi, cat# 80166). After an animal was placed in the channel sled, tubing was connected on both sides, and recording was started. GSH was added using a 1-ml syringe attached to one tube after 2-3min, depending on the behavior of the animal, and recording lasted for approximately 10min. GSH was only added when the animal stayed elongated and did not show contraction or somersaulting behavior. Imaging was performed using the Axio Vert. A1 (Zeiss) with the Colibri 7 as a light source (Zeiss) equipped with the fluorescence filter 38 HE (Zeiss), 5x and 10x Plan Apo objective, and the Axiocam 705 mono (Zeiss). Acquired videos were further processed with Zen Blue 3.4 (Zeiss) to 700x600px, 8-bit and aligned with the Fiji plugin Linear Stack Alignment with SIFT^80^. The aligned stacks were than used for tracing neurons as described by Lagach et al.^81^. Neurons were automatically traced in icy^78^ using the protocol “Detection and Tracking of neurons with emc2”^81^ with individually adjusted parameters depending on the population, magnification, and animal size. Afterwards the quality of the tracks was controlled, and missing links were manually added, or false tracks were removed. For N6 and N4 neurons, tracks were manually separated for the different neuronal sensory (-like) and ganglion.

### GCaMP6S trace analysis

The raw traces were normalized to obtain the fluorescence change ΔF/F0 using the background fluorescence as F0. This background fluorescence was taken by selecting a frame without visible neuronal activity drawing the outline of the animal’s body column and calculating the mean grey therein via Fiji. For further analysis the mean activity of each population or neuronal type was taken with the standard deviation to the mean since it summarized all major events (suppl. Fig. 5). All visualization and normalization were done using customized scripts in R^75^. N3 and N4 spiking frequency was computed using either CASCADE^82^ and/or MATLAB’s (Mathworks) “findpeaks” function with manually adjusted parameters. In all experiments at least 4 animals were used. In Figure 3 A-C only, representative polyps were shown and the mean of the whole neuronal population with the standard deviation, for N4 and N6 divided into sensory and ganglion neurons. More replicates shown in the suppl. Fig. 2-4.

The time sequence of activation of the nerve subpopulation before an opening of the mouth was determined by the time difference between the first activation of the first cell and the opening of the mouth. As the timepoint of first neuronal response, the first activation of the first single cell was taken (shown in Fig. 5G). Higher values respond to an earlier response to GSH. At least 4 animals pre transgenic line were taken.

To find a difference in N6 between germ-free and monocolonized with *Curvibacter* or wildtype microbiota, the area under the curve (AUC) was compared. For this purpose, the mean value of the wildtype microbiota AUC was taken and the difference to the other treatments was calculated. At least 8 animals per treatment were used.

### GCaMP6S and mouth width analysis

For calculation of positive or negative correlations between the mouth opening and the mean activity of the different neuronal subpopulations, the smoothed mouth width data were used. The visualization was done using R and the tidyverse package (Fig 2A-C) ^75,77^. The mouth opening width was adjusted to the scale of the fluorescence change as stated on the right y-axis title. To perform linear correlation analysis, for N6 we compared the fluorescence changes and in case of N3 and N4 the frequency to the mouth opening width at the given time point. For N6, the GCaMP6S traces were divided into ganglion and sensory neurons. Since a continuous signal increase rather than a spiking pattern was observed in N6, we selected a time window of ±15 sec around the mouth opening event and compared this to the mouth width change in the same time window. For N4 and N3 we took the spiking frequency and the width of the mouth opening prior the GSH stimulus and post GSH stimulus. Since we observed a dynamic in the mouth width while recording, we took the minimal mouth width after the mouth opened and the frequency around that time point (30sec window). At least 6 animals per transgenic line were used.

### RNA sequencing and analysis

Transcriptional analysis of *Curvibacter sp.* AEP1.3 was performed by RNA sequencing of bacteria in association with their host and when cultured in R2A without the host (Neogen, cat#NCM0188A). For the latter, 4mL of culture was collected before stationary phase was reached, at an OD_600_ of 0.2-0.3, and centrifuged (4°C, 12000xg). For samples from host-associated *Curvibacter*, 5x500 mono-colonized polyps were prepared as described previously^83^. *Curvibacter* was washed off these animals with PBS and the supernatant was collected and centrifuged (4°C, 12000xg). The bacterial pellet was dissolved in 750µL Trizol by vortexing and 250µL of chloroform was added and samples were centrifuged (12.000xg at 4°C). The aqueous phase was collected and 400µL of 99.9% ethanol was added. The solution was then transferred to silica columns of the ambion PureLink™ RNA Mini Kit (Thermo scientific). RNA was eluted with 35μL RNase free water and stored at −80°C until samples were submitted for sequencing.

Prior to sequencing isolated RNA was treated using the TruSeq stranded total RNA kit (Illumina) and Ribo-Zero Plus kit (Illumina). The remaining RNA was paired end sequenced using a NovaSeq 6000 (Illumina) with 2×150Lbp. RNA sequences were analyzed using the platform Galaxy^84^. The sequences were trimmed using CutAdapt^85^ and Trimmomatic^86^, and MultiQC for quality control^87^. We aligned the reads against the public available *Curvibacter sp.* AEP1.3 genome (ASM216371v1)^83^using Bowtie2^88^. Reads were counted with featureCounts^89^. The normalization of reads and differential gene expression analysis was done using the DeSeq2 pipeline in R^75,90^ and data were visualized using tidyverse in R^77^. All raw RNA-sequence read counts and analyzed data can be found in supplement table 2.

### Statistics

All statistics were done using R and R-studio as IDE^75,76^. In all cases data were tested for their equal variance using Levene’s test and their normal distribution using Shapiro test. Depending on the outcome of those tests either a parametric (t-test, ANOVA, Turkey test) or non-parametric test (Kruskal-Wallis, (pairwise-) Wilcox test, Dunn test) were used. Correction for multiple testing was done using Bonferroni. The replicate number (n) for each dataset is indicated in the figure legends, along with the statistical method used for each comparison and the p value. The cutoff for a significant difference was set as an α < 0.05. Throughout the text, values are reported as median ± standard deviation.

## Supplemental Material

