## Supplementary Material for "Microbes as part of ancestral neuronal circuits: Bacterial produced signals affect neurons controlling eating behavior in *Hydra*"

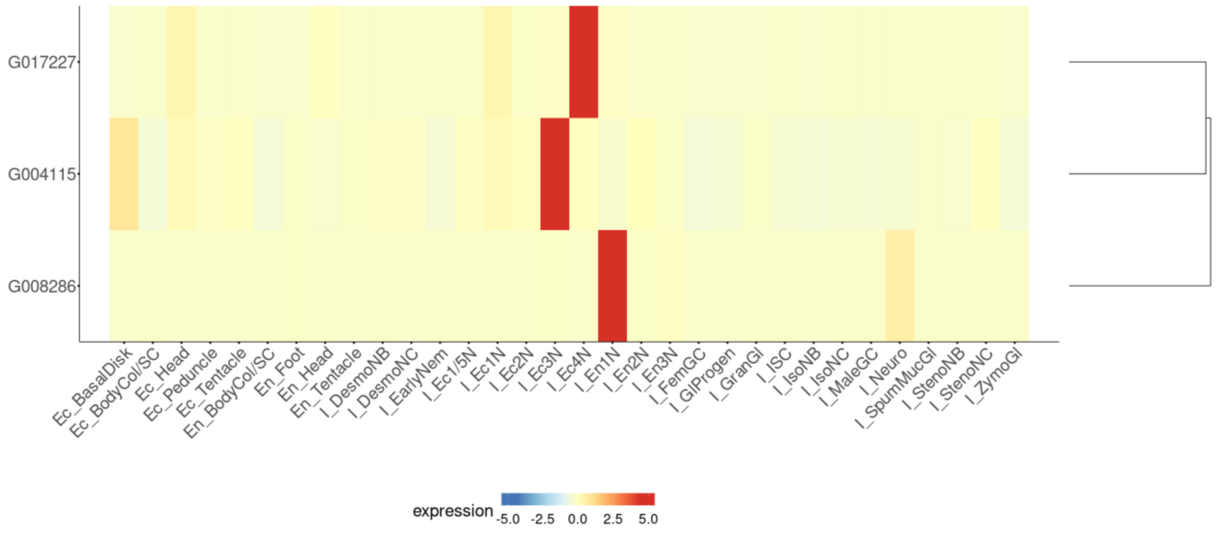


**Supplement Figure 1.** Expression of marker genes used for generating constructs shown in newest single cell atlas with their nomenclature[1]. N6 (i_Ec4N) solely expresses G017227 (t2059aep). N3 (i_Ec3N) solely expresses G004115 (t12874aep). N4 (i_En1N) solely expresses G008286 (t14976aep).

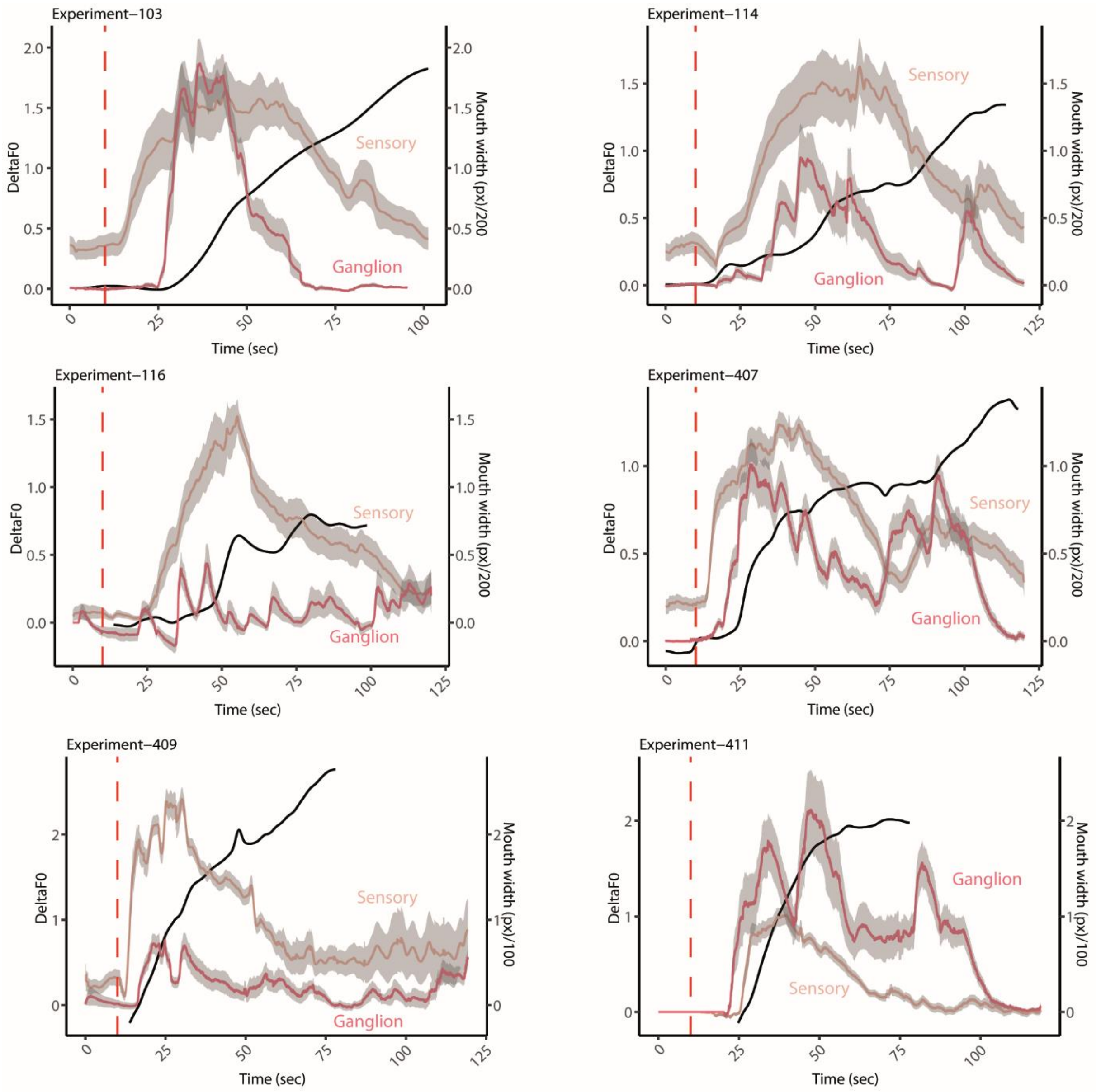


**Supplement Figure 2.** N6 subpopulation activated by GSH shown with GCaMP6S. Here shown multiple animals undergoing the eating behavior. Calcium traces were split into sensory and ganglion neurons. The mean of each cell type population is shown with the standard deviation. In addition, the mouth opening is shown as mouth width over time (measured in pixel) adjusted to the fluorescence change.

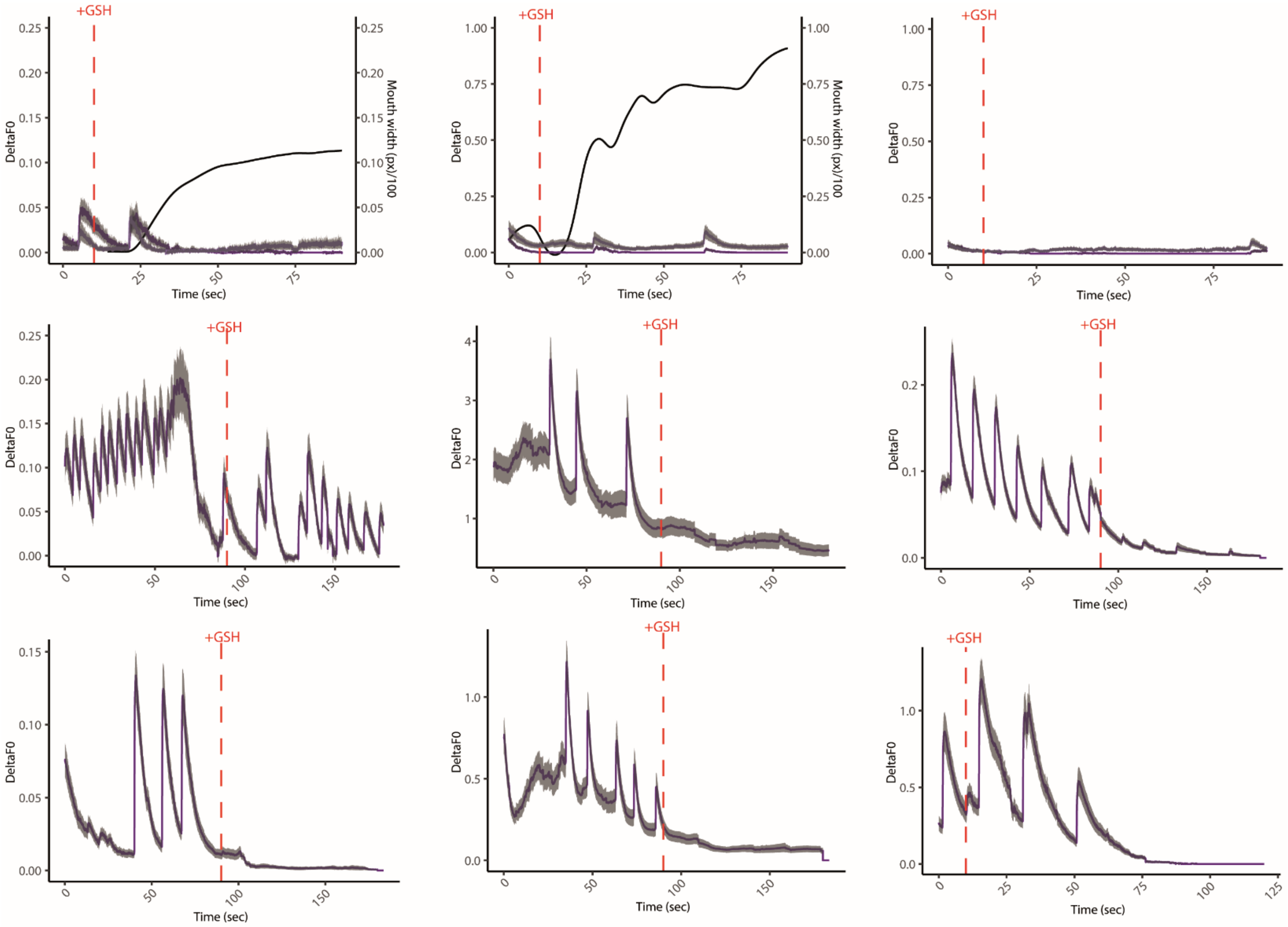


**Supplement Figure 3.** N3 subpopulation activated by GSH shown with GCaMP6S. Here shown multiple animals undergoing the eating behavior. Calcium traces were split into foot and head neurons in the first row otherwise the response of the whole population is shown. The mean of each cell type population is shown with the standard deviation. In addition, the mouth opening is included as mouth width over time (measured in pixel) adjusted to the fluorescence change in the first two plots.

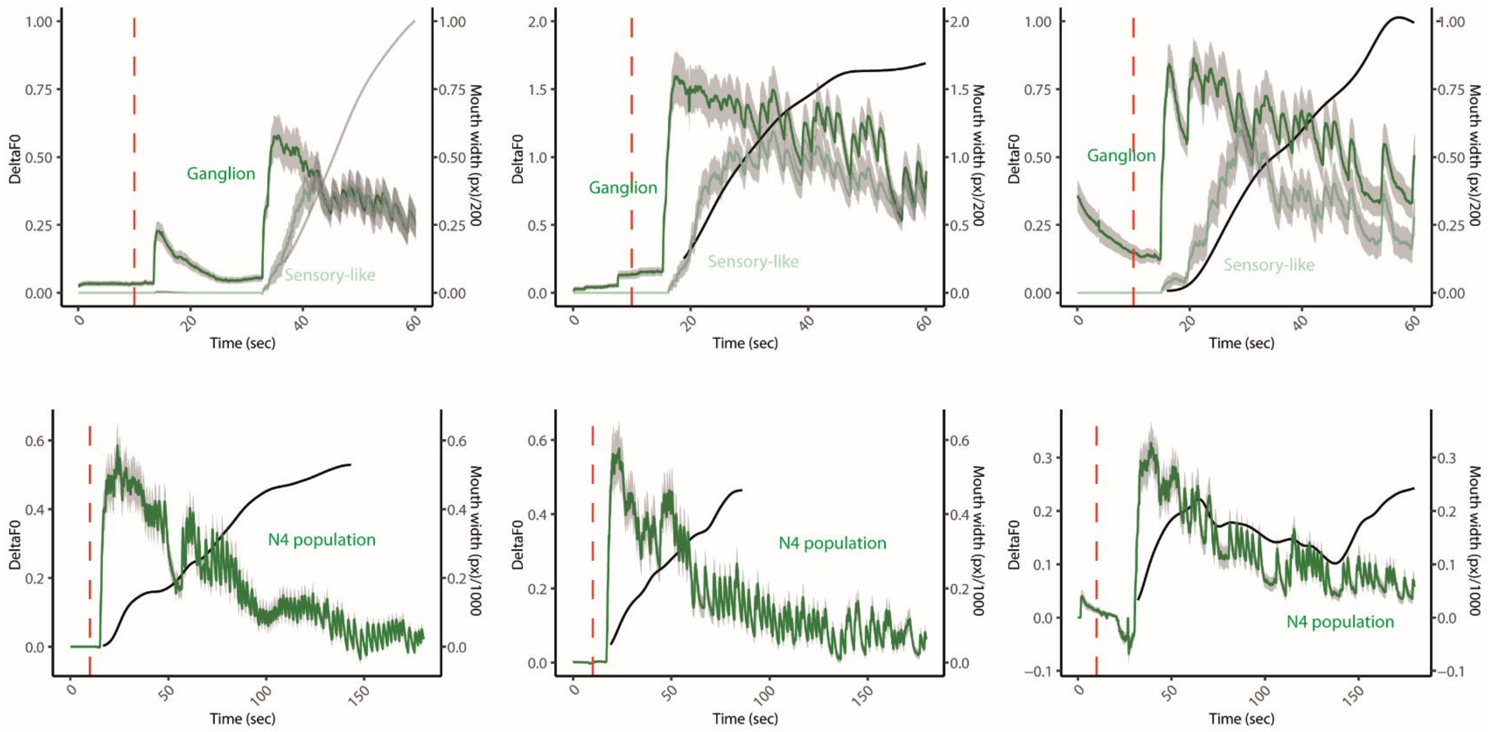


**Supplement Figure 4.** N4 subpopulation activated by GSH shown with GCaMP6S. Here shown multiple animals undergoing the eating behavior. Calcium traces were split into sensory-like and ganglion neurons in the first row otherwise the response of the whole population is shown (mainly ganglion). The mean of each cell type population is shown with the standard deviation. In addition, the mouth opening is shown as mouth width over time (measured in pixel) adjusted to the fluorescence change.

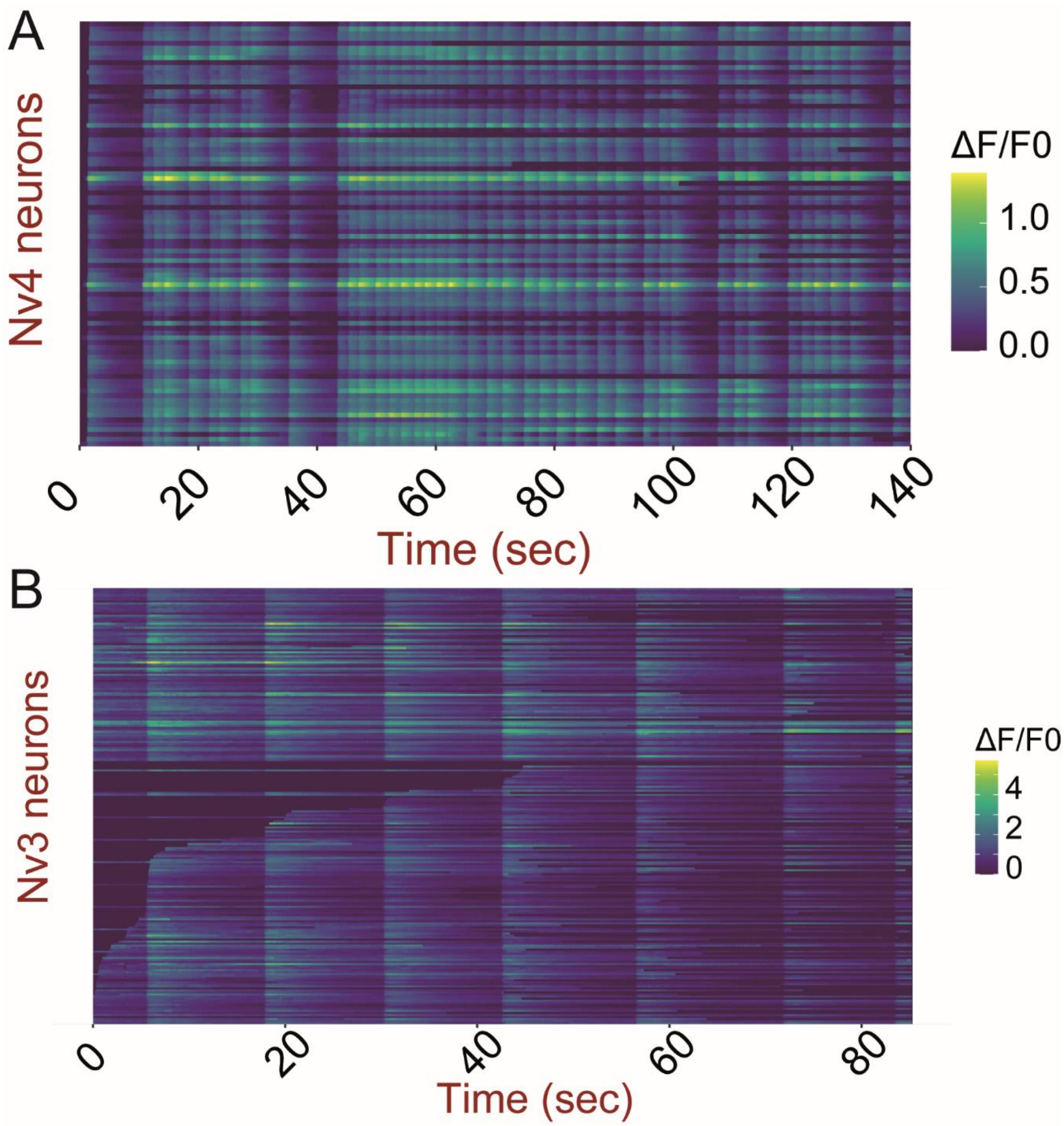


**Supplement Figure 5.** N3 and N4 synchronous firing. Neuronal populations N4 (**A**) and N3 (**B**) show synchronous firing activity in the whole population. **A.** Activity of N4 during the eating behavior. **B.** N3 during an activity episode before addition of GSH.

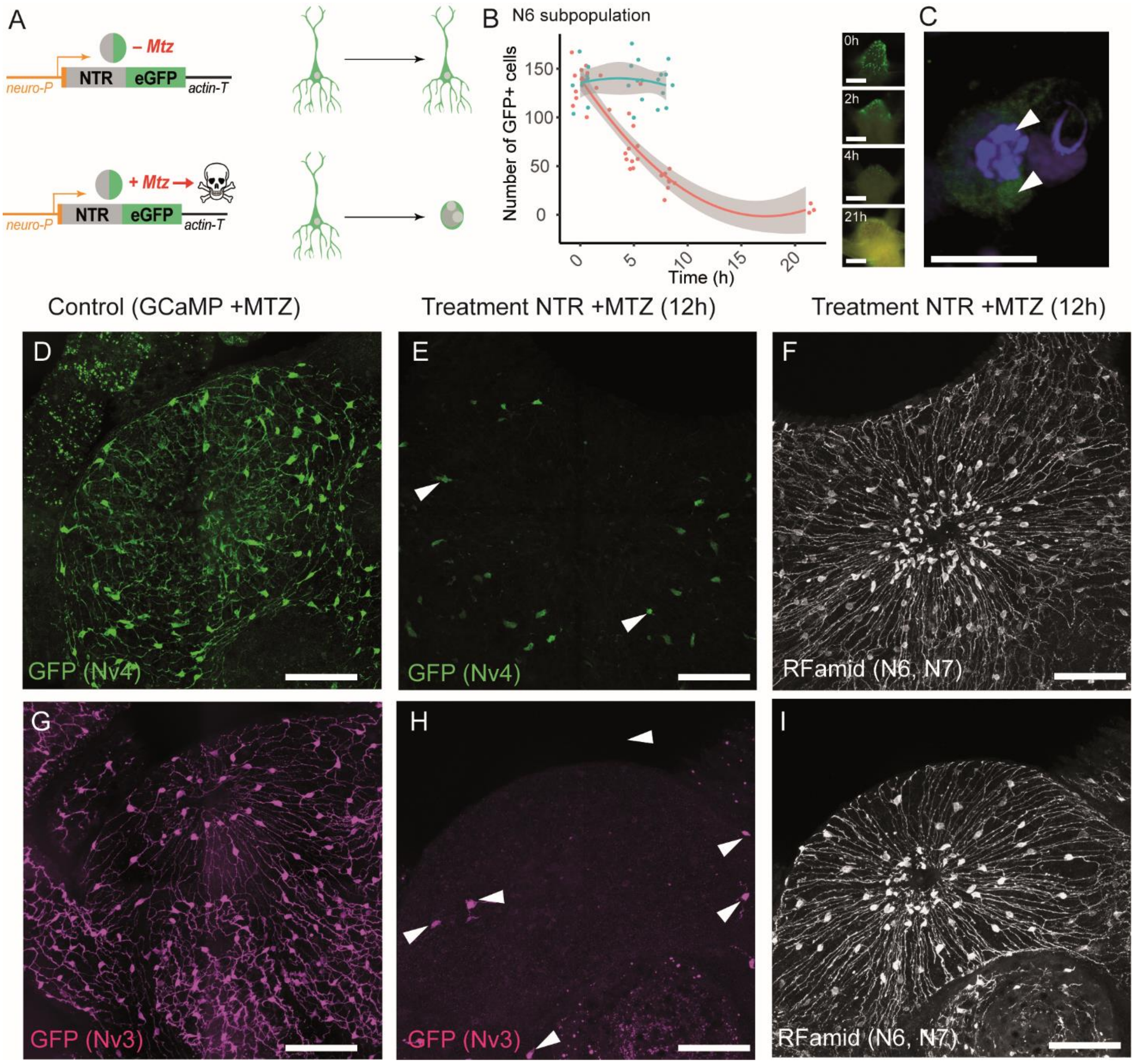


**Supplement Figure 6.** Cell ablation experiment. **A.** Schematic representation of the functioning of the NTR-MTZ system. **B.** Quantification of N6 GFP positive cells *in vivo* after 0h, 5h, 8h and 21h. After 8h till 21h almost all cells were lost. (Scale bar 200µm). **C.** Apoptotic GFP$^{+}$ cell (Scale bar 10µm). **D-F.** Analysis of cell ablation of N4 neurons using antibody staining against GFP and RFamid. **D.** N4::GCaMP6S line with 10mM MTZ after >12h of incubation. **E-F.** N4::NTR-GFP with 10mM MTZ after >12h. **E.** Almost all GFP cells disappeared. **F.** Other neuronal populations are not affected. Here shown by RFamid positive cells (N6 and N7). **G-I.** Analysis of cell ablation of N3 neurons using antibody staining against GFP and RFamid. **D.** N3::GCaMP6S line with 10mM MTZ after >12h of incubation. **E-F.**

N3::NTR-GFP with 10mM MTZ after >12h. **E.** Almost all GFP cells disappeared. **F.** Other populations are not affected. Here shown by RFamid positive cells (N6 and N7). Scale 100µm

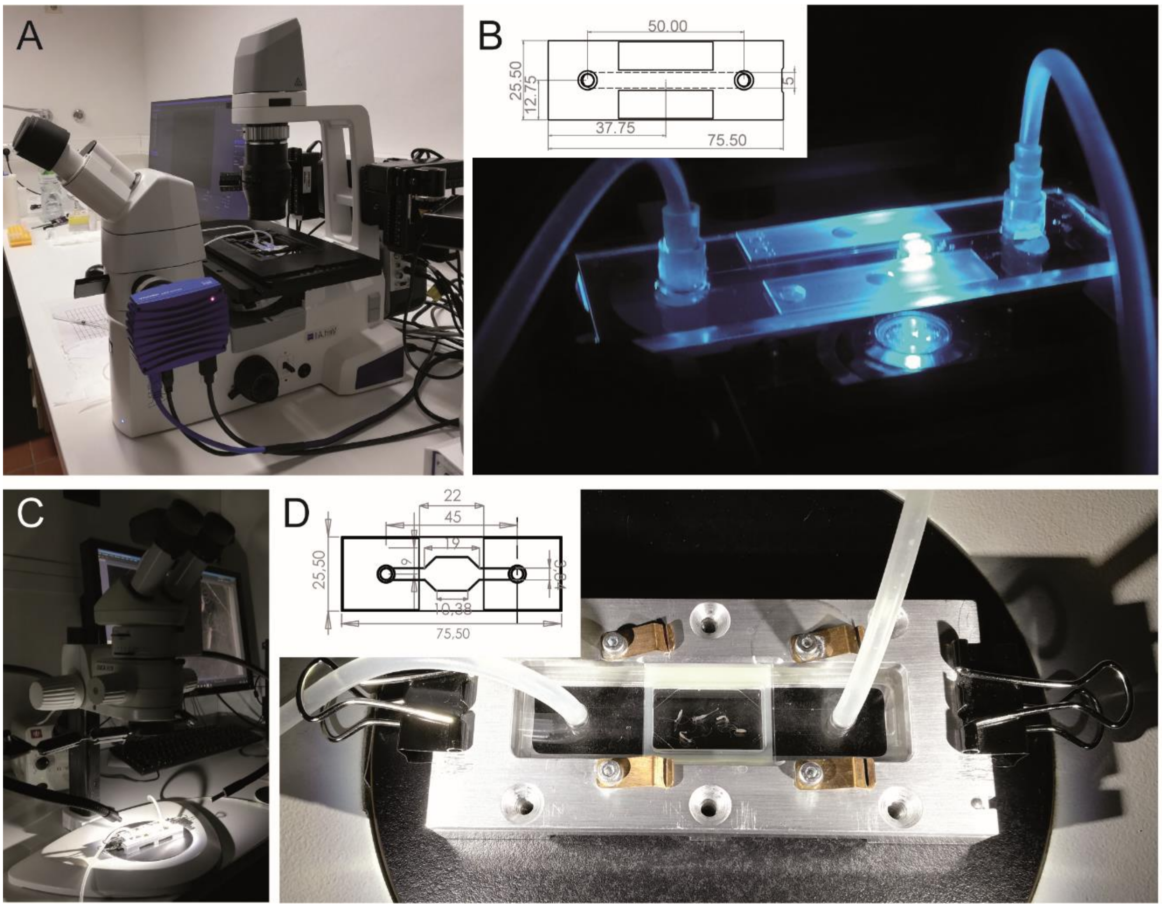


**Supplement Figure 7.** Set up of recording animals for GCaMP6S and animal behavior. **A-B.** Experimental set up for GCaMP6S recordings. **A.** The inverse Zeiss microscope (Axio vert. A1, Zeiss). **B.** The chamber where animals were kept during recording with a technical drawing of the chamber (units in mm, Ibidi cat#80166). **C-D.** Experimental set-up for behavior analysis. **D.** The customized chamber for recording the behavior in the ablation and bacterial manipulation experiments. Technical drawing shown with units in mm.

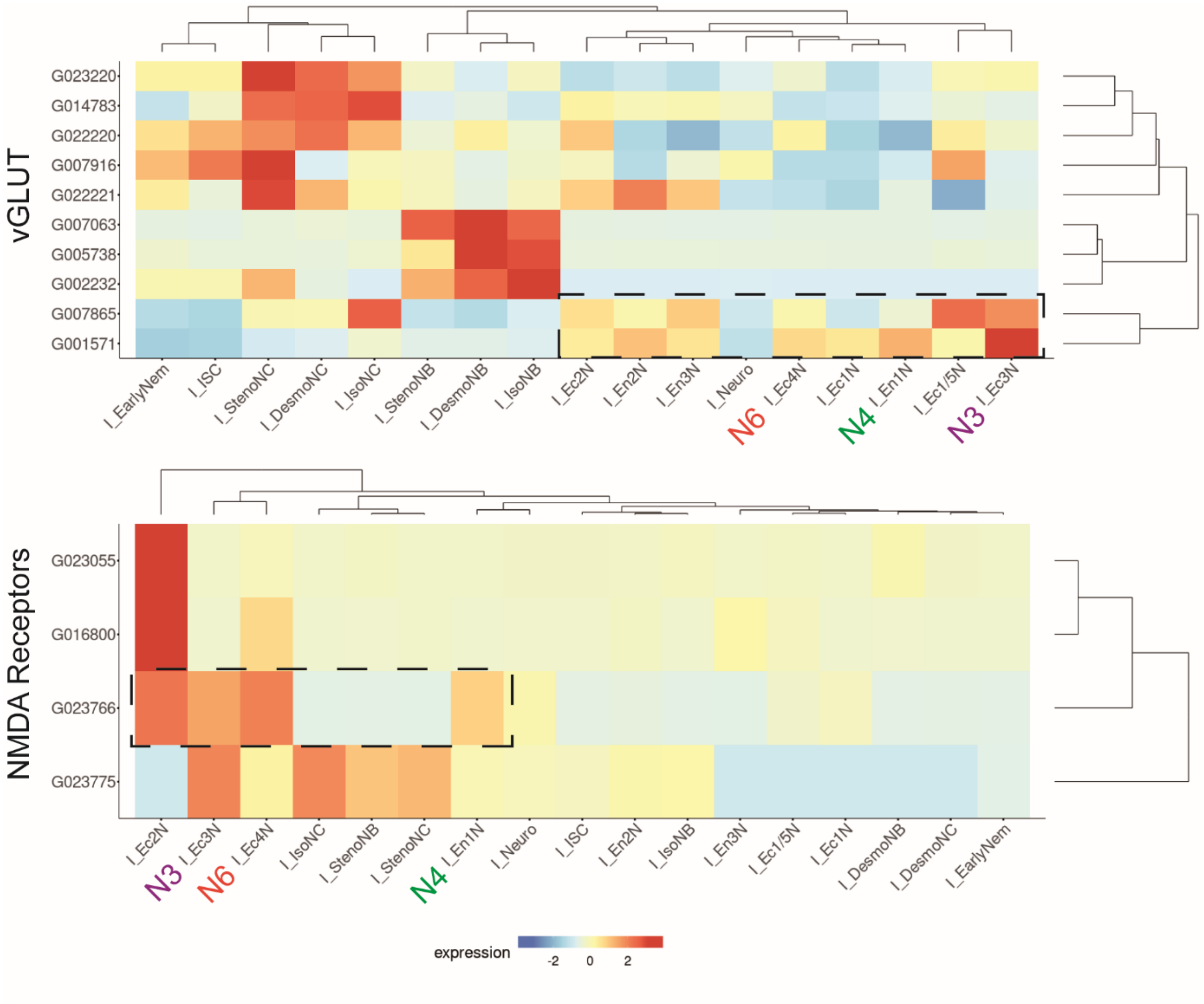


**Supplement Figure 8.** Glutamate (ionotropic) receptor (NMDAR) and transporter in *Hydra's* i-cell lineage (except gland cells). **vGLUT**s are mainly expressed in nematocytes (NC and NB). G007865 and G001571 are expressed mainly in neurons and the strongest in N3 (i_EC3N). Closest sequence to a **NMDA receptor** is expressed in N3, N4 and N6 (G023766, human to hydra: e-value= 0.00). Here different nomenclature was used based on the paper by Cazet et al. 2023.

**Supplement Table 1.** Construct sequences and lines.

**Supplement Table 2.** RNA Sequencing Raw reads raw, analyzed and annotated, related to Figure 6

**Supplement Video1: Behavior annotation, related to all behavioral analysis**

**Supplement Video2: Mouth opening without body and tentacles**

**Supplement Video3: N6 response to GSH 5xObjective, related to Figure 3**

**Supplement Video4: N6 response to GSH 10xObjective, related to Figure 3**

**Supplement Video5: N3 response to GSH 2xObjective, related to Figure 3**

**Supplement Video6: N3 response to GSH 10xObjective, related to Figure 3**

**Supplement Video7: N4 response to GSH 5xObjective, related to Figure 3**

**Supplement Video8: N4 response to GSH 10xObjective, related to Figure 3**
